# Arylsulfatase I is a novel lysosomal chondroitin sulfatase regulating endochondral ossification

**DOI:** 10.1101/2025.10.08.680342

**Authors:** Rafaela Grecco-Machado, Satomi Nadanaka, Yuko Naito-Matsui, Maria Iavazzo, Devin S. Brown, Marziyeh Hassanzadeh, Tuanjie Chang, Elena Polishchuk, Ingrid J. Pickering, Graham N. George, Mark J. Hackett, Nicola Volpi, Carmine Settembre, Hiroshi Kitagawa, B. Frank Eames

**Author notes:** **Corresponding Author**: 2D01–107 Wiggins Road, Saskatoon, SK, S7N5E5, Canada.

## Abstract

During endochondral ossification, chondrocytes undergo maturation and biochemically modify the collagenous extracellular matrix of cartilage. Similar modifications to cartilage proteoglycans (PGs), which are predominantly chondroitin sulfate, have not been characterized. Using synchrotron X-ray fluorescence imaging, we demonstrated that PG sulfation significantly decreased during cartilage maturation of chick embryos. Laser-capture microdissection and RNAseq revealed upregulation of Arylsulfatase I (*Arsi*) in mature cartilage of mouse. ARSI protein also increased in mature cartilage of mouse and chick *in vivo* and during maturation of ATDC5 chondrocytes *in vitro*, whereas expression of the two known chondroitin sulfate PG sulfatases (ARSB and GALNS) was not specific to mature cartilage. Colocalization studies suggested that ARSI is lysosomal, and functional assays revealed that ARSI impacts lysosome homeostasis in chondrocytes. Biochemical analyses of ARSI gain and loss of function cell lines and isolated cell-free systems revealed that ARSI is a novel chondroitin sulfatase, specifically desulfating GalNAc4S at the nonreducing terminal of CS/DS. Finally, *Arsi* knockout in RCS chondrocytes caused increased expression of maturation genes, such as *Col10a1* and *Mmp13*. In total, these data identify ARSI as a novel PG sulfatase regulating endochondral ossification.

## Introduction

During endochondral ossification of the growth plate, mesenchymal cells differentiate into chondrocytes and secrete an extracellular matrix (ECM) of collagens and proteoglycans (PGs) that serves as a cartilage template for subsequent bone formation (Galea et al., 2021). Some of these chondrocytes undergo a developmental transition termed maturation, becoming hypertrophic and biochemically remodelling their collagenous ECM. While the initial cartilage template expresses abundant collagen type 2a1 (*Col2a1*), mature chondrocytes downregulate *Col2a1* expression and upregulate collagen type 10a1 (*Col10a1*; Eames et al., 2003). Degradation of ECM molecules also occurs during cartilage maturation, because mature chondrocytes express collagen-degrading enzymes, such as matrix metalloproteinase 13 (Ortega et al., 2003). Biochemical modifications of PGs remain to be evaluated fully, but PG sulfation might be reduced during cartilage maturation (Deutsch et al., 1995; Farquharson et al., 1994; Hackett et al., 2016; Wuthier, 1969).

While the PG synthesis pathway is well-known, several genes likely involved in reducing PG sulfation remain to be characterized. After PG core proteins are synthesized, a linker tetrasaccharide is added, and then repeating disaccharide glycosaminoglycan (GAG) side chains extend from the linker (Knudson and Knudson, 2001). The repeating disaccharide of heparan sulfate GAGs is glucuronic acid (GlcA)/iduronic acid (IdoA) and *N*-acetyl glucosamine (GlcNAc), while chondroitin sulfate GAGs contain GlcA and *N*-acetyl galactosamine (GalNAc; Brown and Eames, 2016). Cartilage is predominantly comprised of chondroitin sulfate PGs (CSPGs; Gray and Williams, 1989). At this point in their synthesis, PGs get biochemically modified by the addition of *O*–linked sulfate esters at specific GAG residues. CSPGs may be sulfated on the 2-carbon of GlcA and the 4- and 6-carbons of GalNAc (Kusche-Gullberg and Kjellén, 2003). Specific sulfotransferases increase PG sulfation, and specific sulfatases reduce PG sulfation (Sardiello et al., 2005). Among 17 sulfatases that have been identified in the human genome from gene sequence analyses (Sardiello et al., 2005), only two are known to specifically desulfate CSPGs. Arylsulfatase b (ARSB) acts on 4-sulfated chondroitin (C4S), and GALNS acts on C6S (Brown and Eames, 2016). Much must be learned about sulfatases, because the specific biochemical substrates of six putative sulfatases, including ARSI, are not known (Schlotawa et al., 2020).

PG sulfatases can be related functionally to lysosomes. Lysosomes are membrane-bound degradative organelles that contain hydrolytic enzymes that break down macromolecules such as proteins, lipids, carbohydrates, and nucleic acids into smaller components that can be reused (Settembre and Perera, 2024). Many sulfatases, such as ARSB, GNS, IDS, and GALNS, colocalize with lysosomes and play key roles in degradation of PGs (Brown and Eames, 2016). ARSI has no known cellular localization (Schlotawa et al., 2020). A relationship between sulfatase function and lysosome biogenesis has been established. Genetic mutations in lysosomal sulfatases lead to inherited metabolic diseases known as mucopolysaccharidosis (MPS), characterized by lysosomal accumulation (storage) of undegraded PG molecules (Hers, 1965). Cellular homeostasis of lysosomes, including numbers and sizes, is altered in MPS ( Bajaj et al., 2019; Settembre and Perera, 2024).

PG sulfation influences cartilage function and can affect endochondral ossification (Dzobo et al., 2016; Gama et al., 2006; Gao et al., 2014). The negative charge of PG sulfation attracts more water, increasing the compressive-resistant strength of cartilage (Chahine et al., 2005). Inadequate PG sulfation can cause a variety of skeletal defects during development. Mice with mutations in either *Papss2* (regulates the sulfate donor within the cell)*, Slc26a2* (transports sulfate within the cell), or *Sumf1* (activates sulfatases) have reduced chondrocyte proliferation, severe growth retardation, and delayed secondary ossification center formation (Cortes et al., 2009; Gualeni et al., 2010; Settembre et al., 2007; Settembre et al., 2008). In human patients, mutations to *ARSB* or *GALNS* can cause MPS types IV or VI, respectively (Muenzer, 2011). Both MPS IV and VI cause reduced growth of bones that form by endochondral ossification, but the underlying cellular and molecular mechanisms are currently unknown (Montano et al., 2007; Valayannopoulos et al., 2010).

Given the relevance of PG sulfatases to human disease, the biochemical functions of the remaining orphan sulfatases need to be revealed. Initially, we used synchrotron-based X-ray fluorescence (XRF) imaging to reveal that PG sulfation significantly decreases during cartilage maturation. Supporting the hypothesis that a chondroitin sulfatase was expressed during cartilage maturation, laser-capture microdissection/RNAseq and other studies demonstrated that expression of ARSI, but not ARSB nor GALNS, increased as sulfate esters decreased in developing mature cartilage. Gain- and loss-of-function studies *in vitro* revealed that ARSI is a novel, lysosomal chondroitin sulfatase that regulates lysosome homeostasis and maturation genes in chondrocytes, opening the pathway for understanding how biochemical modification of PGs affects endochondral ossification.

## Results

### PG sulfation decreased during cartilage maturation

Levels of PG sulfation were measured during cartilage maturation of the HH36 chick humerus, which undergoes endochondral ossification. Mature cartilage regions were identified using histological and molecular analyses. Whole-mount Alizarin red staining, which binds mineralized tissue, revealed perichondral bone in the mid-diaphysis (Fig. 1A). Whole-mount staining of Alcian blue, which binds sulfated PGs, appeared reduced in the cartilage region that is directly underneath perichondral bone (Fig. 1B). Similarly, reduced staining levels of Safranin O, which also binds sulfated PGs, on histological sections of the HH36 chick humerus were apparent in this cartilage region, which was confirmed as mature cartilage by the presence of hypertrophic chondrocytes and expression of Collagen type X (COLX; Fig. 1C’,C’’,D; Eames et al., 2003).

**Figure 1.**
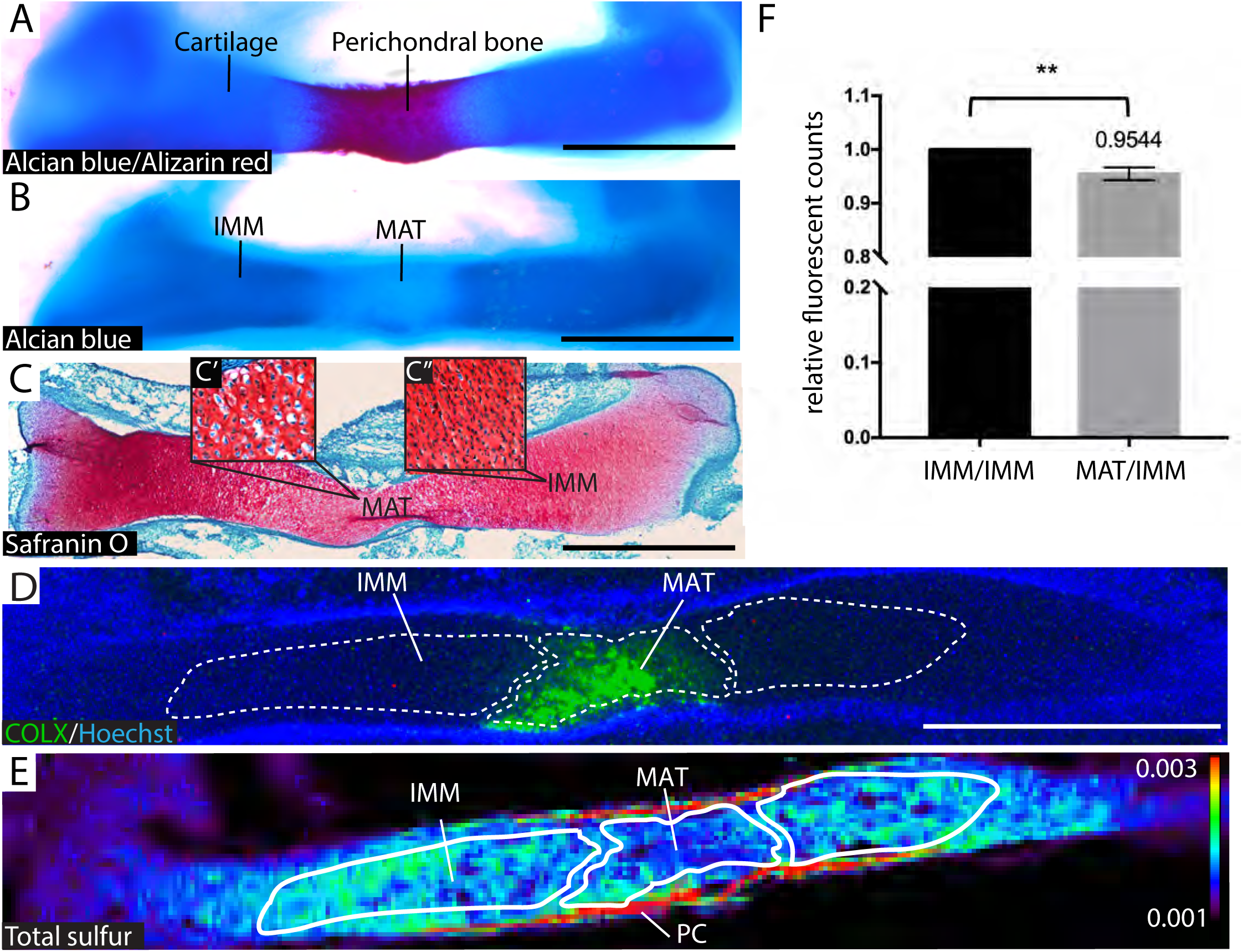
Sulfation decreased during cartilage maturation of the HH36 chick humerus. **(A)** Whole-mount Alcian blue and Alizarin red staining indicated cartilage and perichondral bone regions, respectively. **(B)** Whole-mount Alcian blue appeared decreased in mature cartilage (MAT) underlying perichondral bone, compared to immature cartilage (IMM). **(C)** Similarly, Safranin O staining of sections appeared decreased in mature cartilage and showed hypertrophic chondrocytes in MAT (C’) and non-hypertrophic chondrocytes in IMM (C’’). **(D)** COLX expression verified MAT on sections and was used to define MAT and IMM regions (white dashed lines) for XRF image analysis. **(E)** XRF imaging of adjacent sections suggested that total sulfur levels decreased in MAT. **(F)** Quantitation of XRF images (n=9) and one-sample t-test revealed a significant decrease of 4.5 ± 1.2% in the average total sulfur in MAT, relative to IMM. **p<0.005. Abbreviations: IMM=immature cartilage; MAT=mature cartilage; PC=perichondrium. Scale bars: A-D=1mm.

Synchrotron-based XRF imaging of sections of the HH36 chick humerus revealed that sulfur was enriched in cartilage, relative to surrounding tissues (n=9; Fig. 1E), as previously reported (Hackett et al., 2016). To quantitate and compare each cartilage region’s sulfur levels, COLX immunostaining on adjacent sections was used to define mature cartilage, as well as nearby immature cartilage (Fig. 1D, dotted lines). Qualitatively, COLX immunostained regions mostly fit well within a low sulfur region (Fig. 1E). Quantitation of sulfur levels in these identified regions verified a significant decrease (4.5 ± 1.2%) in the average total sulfur content in mature cartilage compared to immature cartilage (Fig. 1F). The resting and proliferative zones in the extreme epiphyses of developing cartilage also appeared to have lower levels of total sulfur, although this was not quantitated.

A diversity of chemical forms of sulfur are present in biological tissues like cartilage, but PG sulfation is reflected specifically by sulfate esters (Hackett et al., 2016; Mikami and Kitagawa, 2013). To investigate relative levels of specific chemical forms of sulfur in developing immature and mature cartilage, X-ray absorption near edge structure (XANES) analysis was performed on HH36 chick humerus sections. Combined fittings of five sulfur chemical forms from standard curve measures (among eight tested) matched very well the averaged XANES spot scans (n=4; Supplemental Fig. S1). *O*-linked sulfate esters were the most abundant (77.3 ± 2.1% of sulfur in immature cartilage; 69.5% ± 10.7% in mature cartilage), followed by sulfonic acids, thioethers, disulfides, and sulfoxides (Table 1). *N*-linked sulfate esters were practically negligible in developing cartilage and were not included in the final XANES fitting, as reported previously (Hackett et al., 2016). The fact that *O*-linked sulfate esters were the most abundant form of sulfur detected by XANES in developing chick cartilage suggested that the majority of sulfur detected by XRF was from PGs.

**Table 1.**
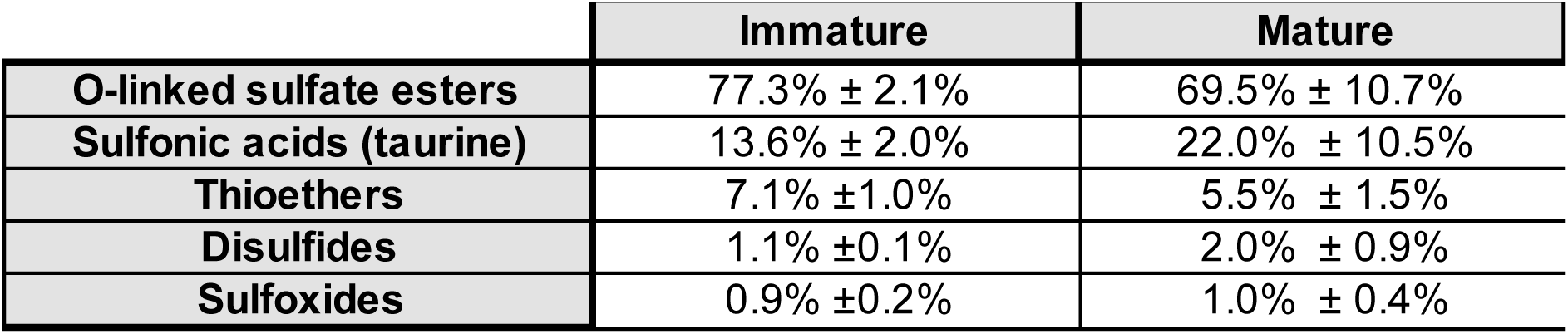
XANES measures revealed *O*-linked sulfate esters were the majority of sulfur forms in the HH36 chick humerus.

To visualize the distribution of different chemical forms of sulfur in developing cartilage *in situ*, chemically specific XRF maps were generated (n=5). COLX immunostaining on adjacent sections was used to define the mature and nearby immature cartilage regions (dotted lines, Fig. 2A). Qualitatively, the relative intensities of total sulfur and sulfate esters had similar distributions in the HH36 chick humerus, with lower levels apparent in the mature cartilage region (Fig. 2B,C). The similarity in maps of total sulfur and sulfate esters detected by chemically-specific XRF supported XANES results that PG sulfation accounted for the majority of sulfur in cartilage. By contrast, maps of sulfonic acid and lower oxidation states of sulfur did not clearly follow this distribution pattern across developing cartilage (Fig. 2D,E). Quantitation of chemically specific XRF maps revealed that sulfate esters decreased significantly (13.4 ± 4.1%) in mature cartilage, compared to immature cartilage (Fig. 2F). Total sulfur also tended to decrease (8.8 ± 4.2%) in mature cartilage, but not significantly in these samples (Fig. 2F). Like the XANES analyses (Table 1), sulfonic acids tended to increase slightly in mature cartilage (25.4 ± 35%), but not significantly, likely due to large variation between samples (Fig. 2F). In summary, significant decreases in *O*-linked sulfate esters demonstrated that PG sulfation decreases during cartilage maturation.

**Figure 2.**
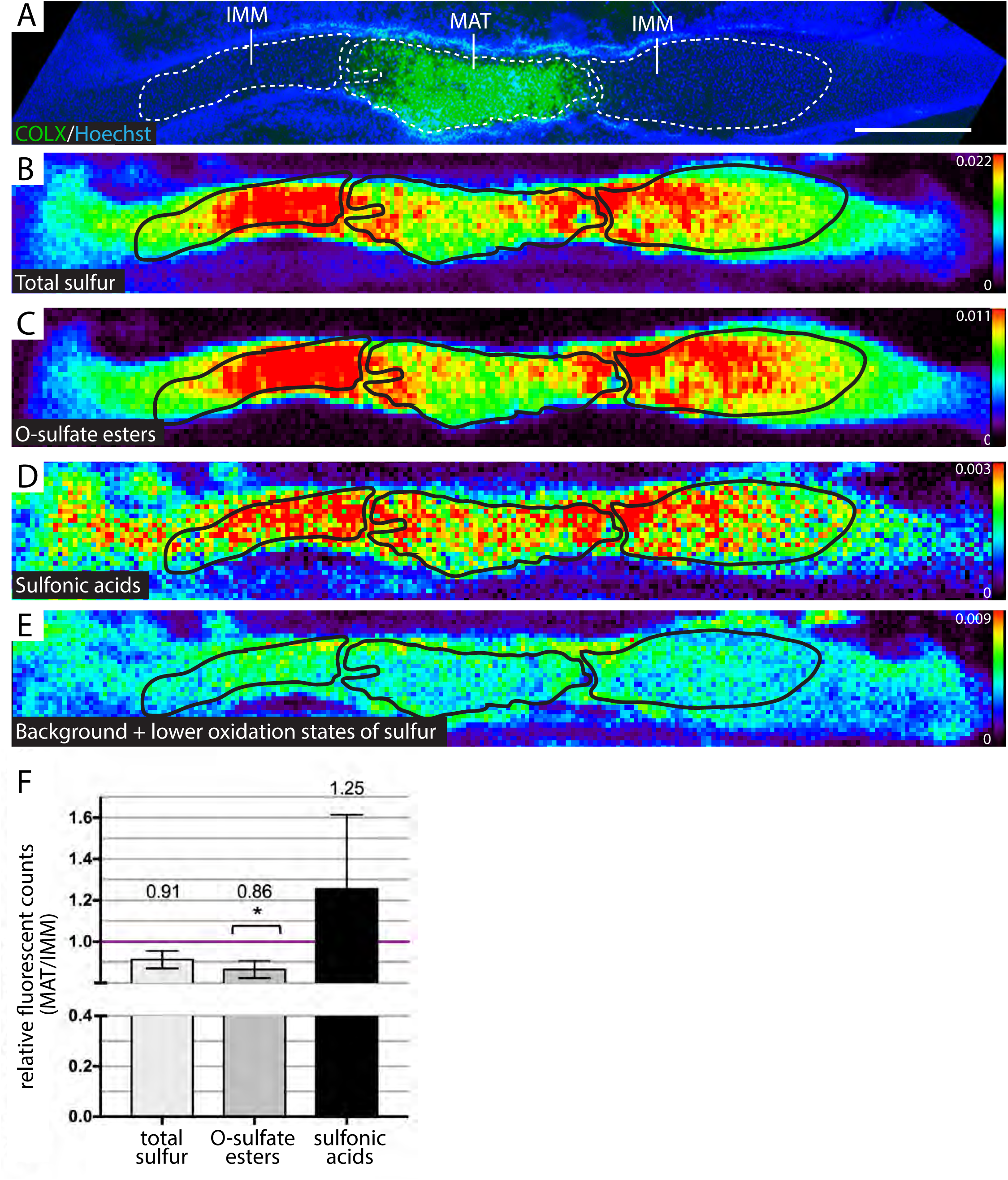
Chemically specific XRF maps of the HH36 chick humerus revealed a significant decrease in *O*-sulfate esters in mature cartilage. (**A**) COLX immunohistochemistry defined mature (MAT) and nearby immature (IMM) cartilage regions of interest (dotted lines). Levels of total sulfur **(B)** and *O*-sulfate esters **(C)** had similar distributions across developing cartilage, with lower levels in MAT. Levels of sulfonic acid **(D)** and lower oxidation states of sulfur **(E)** did not follow a clear distribution pattern across developing cartilage. **(F)** Quantitation of MAT and IMM (black outlines in B-E) on XRF images (n=5) showed that *O*-sulfate esters significantly decreased 13.4 ± 4.1% in MAT. *p<0.003. Abbreviations: IMM=immature cartilage; MAT=mature cartilage. Scale bar=1mm.

Since loss of PG sulfation is linked to PG degradation (Settembre et al., 2007; Settembre et al., 2008), overall loss of PGs during cartilage maturation was investigated using Fourier transform infrared (FTIR) imaging on sections of the HH36 chick humerus (n=7; Supplemental Methods). Using previously-reported wave numbers, proteins (Amide I: 1590-1720cm^-1^; Boskey and Pleshko Camacho, 2007) tended to decrease in mature cartilage (5 ± 3%), compared to immature cartilage, although this was not statistically significant (Supplemental Fig. S2B,E). Since proteins were somewhat decreased, PG and GAG FTIR maps were normalized to the integrated Amide I peak. With or without this normalization, PGs (985-1140cm^-1^; Saarakkala and Julkunen, 2010) and GAGs (second derivative peak 1374cm^-1^; Rieppo et al., 2012) also both tended to decrease in mature cartilage (4 ± 4% and 3 ± 3%, respectively), but not significantly (Supplemental Fig. S2C-E). These FTIR imaging data suggested that at least some of the sulfate esters in mature cartilage decreased independently of overall PG levels.

### ARSI, but not GALNS or ARSB, was expressed specifically in mature cartilage in mouse and chick

To test the hypothesis that the significant decrease in PG sulfation during cartilage maturation was due to a PG sulfatase expressed specifically in mature cartilage, our published transcriptomic dataset (Gomez-Picos et al., 2025) from laser capture microdissected mature and immature cartilage (n=3 for each tissue) in the E14.5 mouse humerus was analyzed. *Galns* and *Arsb* encode the only known animal sulfatases that specifically remove sulfate esters from CSPGs, the main PG in cartilage matrix (Brown and Eames, 2016). However, neither *Galns* nor *Arsb* transcripts were increased in mature cartilage (Table 2). Genes encoding known heparan sulfate PG sulfatases, such as *Gns, Ids, Arsg,* and *Sulf1* (Brown and Eames, 2016), also were not enriched in mature cartilage; *Sulf2* was even significantly downregulated (Table 2). Of the putative PG sulfatases, including *Arsd, Arse, Arsf, Arsh, Arsi,* and *Arsj*, only *Arsi* was significantly increased (1.65 log2FC) in mature over immature cartilage (Table 2). The classic maturation marker *Col10a1* was also enriched (10.62 log2FC) in mature cartilage in these datasets (Table 2). In summary, transcriptomic analyses identified that the gene *Arsi*, encoding an orphan sulfatase, was specifically expressed during cartilage maturation.

**Table 2.**
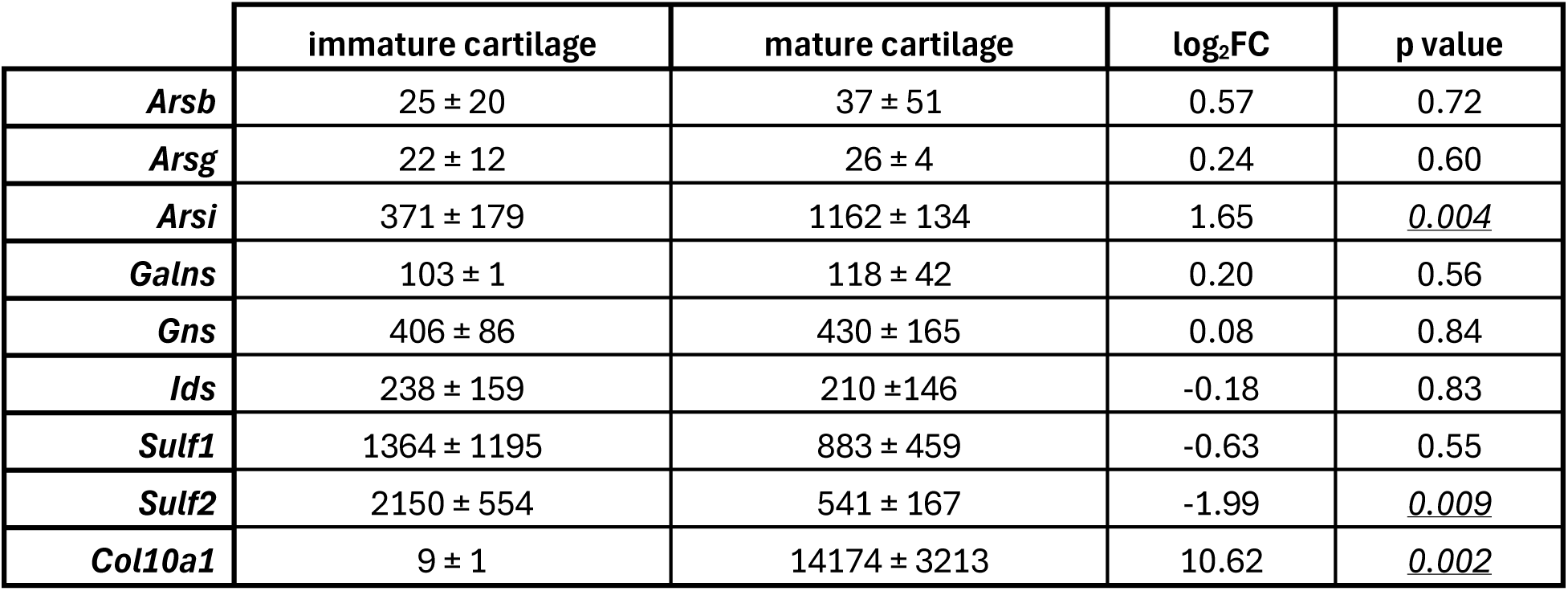
The putative PG sulfatase *ARSI* was the only sulfatase significantly increased in mature cartilage of the E14.5 mouse humerus.

|  | immature cartilage | mature cartilage | log<sub>2</sub>FC | p value |
| --- | --- | --- | --- | --- |
| *Arsb* | 25 ± 20 | 37 ± 51 | 0.57 | 0.72 |
| *Arsg* | 22 ± 12 | 26 ± 4 | 0.24 | 0.60 |
| *Arsi* | 371 ± 179 | 1162 ± 134 | 1.65 | <u>0.004</u> |
| *Galns* | 103 ± 1 | 118 ± 42 | 0.20 | 0.56 |
| *Gns* | 406 ± 86 | 430 ± 165 | 0.08 | 0.84 |
| *Ids* | 238 ± 159 | 210 ± 146 | -0.18 | 0.83 |
| *Sulf1* | 1364 ± 1195 | 883 ± 459 | -0.63 | 0.55 |
| *Sulf2* | 2150 ± 554 | 541 ± 167 | -1.99 | <u>0.009</u> |
| *Col10a1* | 9 ± 1 | 14174 ± 3213 | 10.62 | <u>0.002</u> |

To verify that analyses in different species did not confound these results, ARSI sequence conservation was evaluated, and ARSI expression was analyzed in both chick and mouse. ARSI protein sequence was highly conserved across vertebrates, even more so when comparing just the putative sulfatase domain, suggesting functional conservation (Table 3, Supplemental Fig. S3, Supplemental Table 1). The sulfatase domain of human ARSI was 97.4% similar with *Mus musculus*, 90.0% with *Gallus gallus*, 75.3% with *Danio rerio* Arsia, and 74.7% with *Danio rerio* Arsib (zebrafish retained duplicate copies of ARSI after the teleost genome duplication; Amores et al., 1998).

**Table 3.**
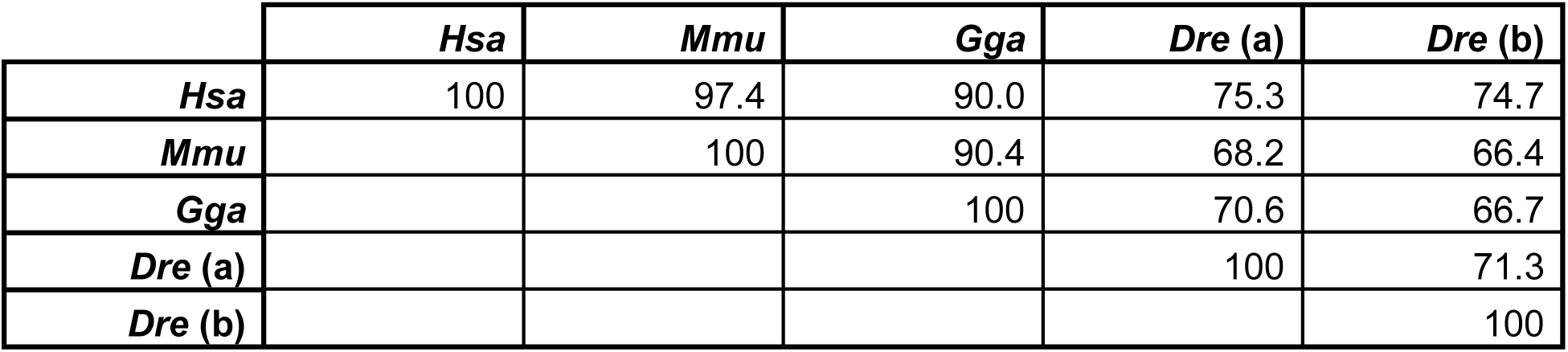
Amino acid sequences of the putative sulfatase domain of ARSI were highly conserved across vertebrate clades. Numbers reflect % identity. Abbreviations: (a)=Arsia; (b)=Arsib; *Dre*=*Danio rerio*; *Gga=Gallus gallus*; *Hsa*=*Homo sapiens*; *Mmu=Mus musculus*.

To analyze ARSI expression in chick cartilage, RNA *in situ* hybridization was performed (Supplemental Methods). *ARSI* mRNA expression was confirmed in mature cartilage of the HH36 chick humerus (n=3), in similar domains as the prehypertrophic marker *IHH*, but restricted to superficial regions of the cartilage (Supplemental Fig. S4A-B’). *ARSI* was not expressed in more mature regions of mature cartilage, such as where *COL10A1* or *SPP1* were expressed, or where *COL2A1* was downregulated (Supplemental Fig. S4C-E’). These expression results were confirmed with two other probes targeting different portions of the *ARSI* mRNA (data not shown). Transcripts for the chondroitin sulfatase genes *GALNS* and *ARSB* did not show any specific expression in immature or mature cartilage domains of the HH36 chick humerus (Supplemental Fig. S4F-G’).

Immunostaining was used to further confirm expression of ARSI in mature cartilage of the chick. In the HH36 chick humerus (n=10), hypertrophic chondrocytes and COLX expression on adjacent sections were again used to estimate the region of mature cartilage (Fig. 3A,B; dotted lines in all panels). ARSI was restricted to mature cartilage, but dual immunostaining with COLX confirmed that ARSI protein was in the hypertrophic domain of mature cartilage (Fig. 3C,D). COLX immunoreactivity was in the extracellular matrix, but ARSI immunoreactivity did not overlap with COLX, appearing closer to nuclei, suggesting intracellular localization (Fig. 3B-D’). Similar to mRNA expression, ARSI protein was expressed in more superficial regions of mature cartilage (Supplemental Fig. S5). Neither of the two known CSPG sulfatases, GALNS and ARSB, showed increased expression in mature cartilage (Fig. 3E,F). GALNS was not above background levels in any region of the humerus. ARSB was expressed at similar levels in both mature and immature cartilage regions, with apparent intracellular immunoreactivity (Fig. 3F’).

**Figure 3.**
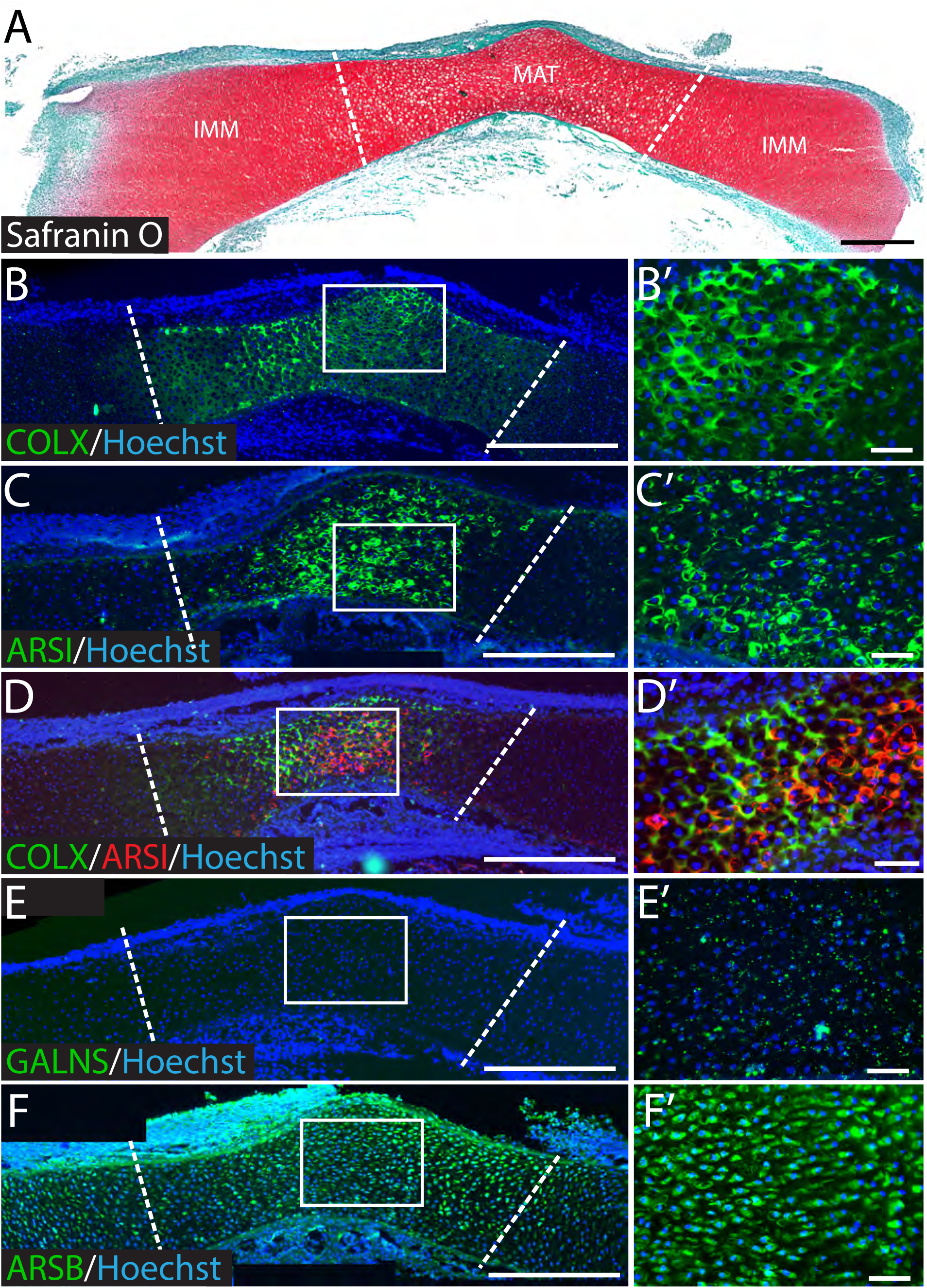
Immunostaining demonstrated that ARSI, but not GALNS or ARSB, was specifically expressed in mature cartilage of the HH36 chick humerus. **(A)** Safranin O staining showed hypertrophic chondrocytes in mature cartilage (dotted lines in all panels estimate the mature cartilage region). In addition to COLX **(B)**, ARSI **(C)** was expressed in mature cartilage (n=10). **(D)** Dual immunostaining for COLX (green) and ARSI (red) confirmed their expression in mature cartilage, but also suggested that ARSI was expressed intracellularly. In both regions of the humerus, GALNS **(E)** was not observed above background levels, while ARSB **(F)** was observed in similar levels. Boxed regions in panels A-F indicate regions shown in panels A’-F’, respectively. Abbreviations: IMM=immature cartilage, MAT=mature cartilage. Scale bars: A-F=400μm; A’-F’=50μm.

To further investigate conservation of ARSI expression during chondrocyte maturation, immunostaining of the E14.5 mouse humerus and a mouse chondrocyte cell line, ATDC5 (Supplemental Methods), were performed. Indeed, ARSI and COLX immunostaining were in similar domains of mature cartilage in the E14.5 mouse humerus (Supplemental Fig. S6A,B). Cartilage differentiation in micromasses of ATDC5 cells was confirmed by Alcian blue staining, and increased COLX immunostaining was apparent during ATDC5 chondrocyte maturation (Supplemental Fig. S6C-H). Immunostaining and Western blot analyses also showed that ARSI levels increased during ATDC5 chondrocyte maturation (Supplemental Fig. S6I-L). In sum, these *in vitro* and *in vivo* protein expression data confirmed transcript data in chick and mouse, demonstrating conserved, specific expression of ARSI in mature cartilage, where XRF imaging showed that PG sulfation was decreased.

### ARSI colocalized with lysosomes and functionally impacted lysosome homeostasis

To confirm immunostaining results suggesting that ARSI was localized intracellularly, and to identify its cellular compartments, imaging was undertaken of cells stably expressing fluorescent-tagged human ARSI. Human ARSI with EGFP fused to its C-terminus was expressed in HeLa cells (Fig. 4A). Dual immunostaining revealed that ARSI was colocalized with the lysosome marker LAMP1 and the endosome marker RAB5 (Fig. 4B-G’; Moss et al., 2020), suggesting that ARSI is a bona fide lysosomal enzyme. Colocalization of ARSI and lysosomes was also demonstrated by expressing human ARSI with mCherry tagged to its C-terminus in ATDC5 mouse chondrocytes and imaging with the vital dye LysoTracker (data not shown).

**Figure 4.**
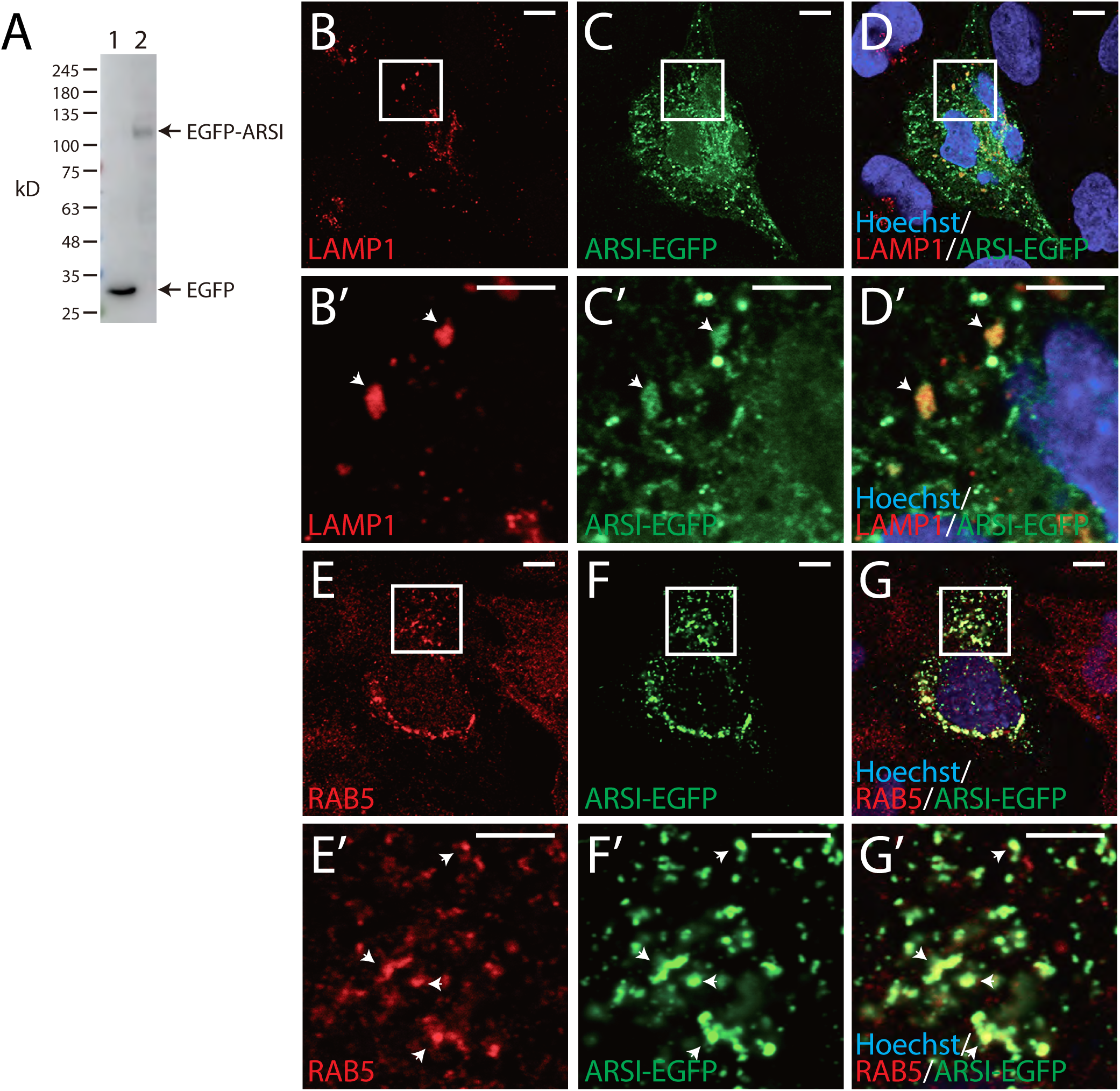
EGFP-tagged ARSI colocalized with lysosomes and endosomes in HeLa cells. **(A)** Anti-GFP Western blot demonstrated expression of EGFP (lane 1) EGFP-tagged human ARSI (Lane 2) in HeLa cells 2 days after transfection. Immunostaining showed colocalization (white arrowheads) of LAMP1 and ARSI **(B-D)** and colocalization of RAB5 and ARSI **(E-G)** in HeLa cells. Anti-GFP was used for immunostaining of ARSI. Boxed regions in panels B-G indicate regions shown in panels B’-G’, respectively. Scale bars: B-G=10μm; B’-G’=5μm.

To evaluate the impact of ARSI on lysosome homeostasis, *Arsi* loss of function was created in the RCS line of rat chondrocytes, and lysosome numbers and sizes were analyzed. CRISPR targetting of *Arsi* was confirmed by Western blotting of two independent clones, both of which had a lack of ARSI protein expression (Fig. 5A). Three independent quantitative measures of lysosomes were performed. FACS analyses of RCS chondrocytes labelled with the vital dye LysoTracker demonstrated that fluorescence intensity was significantly increased in two *Arsi^-/-^* chondrocyte clones, compared to a parental control clone (Fig. 5B). Those two *Arsi^-/-^* clones also had significant increases in lysosome numbers, compared to a parental control, upon quantitation of immunostaining with LAMP1 (Fig. 5C-F). Finally, quantitation in TEM images of one the *Arsi^-/-^* clones showed significantly larger lysosomes, compared to a parental clone (Fig. 5G-I). In total, these data revealed that ARSI is lysosomal and regulates lysosome homeostasis.

**Figure 5.**
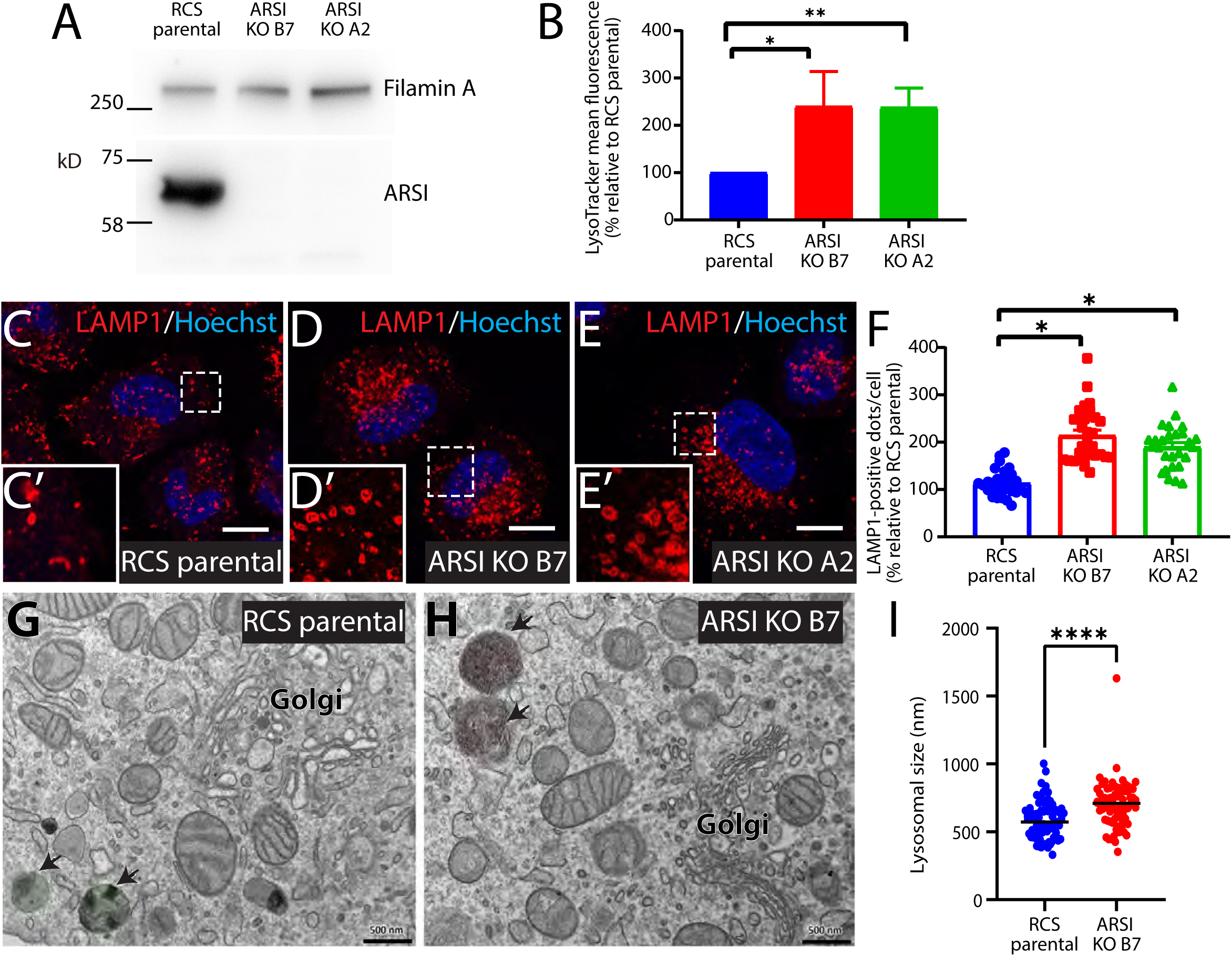
ARSI inhibited lysosome number and size in RCS chondrocytes. **(A)** Western blot confirmed loss of ARSI protein in two CRISPR-targetted RCS chondrocyte clones. Filamin A was used as a loading control. **(B)** FACS analysis labeled with LysoTracker and Hoechst dyes revealed that two ARSI knockout (KO) RCS chondrocyte clones had increased lysosomes per cell, compared to parental control (n=3, each clone). **(C-F)** Immunostaining of LAMP1 (red) with Hoechst dye (blue) in clones of parental (C) and two ARSI knockout (D,E) RCS chondrocytes demonstrated increased lysosomes, which was confirmed by quantification of LAMP1-positive dots/cell (n=3, each clone). Dashed boxed regions in panels C-E indicate regions shown in panels C’-E’. **(G-I)** TEM images of RCS chondrocytes also showed smaller lysosomes in a parental clone (G), compared to an ARSI knockout clone (H), which was verified by quantitation (I). Arrows indicate lysosomes. *p<0.05; **p<0.005; ****p<0.0005. Abbreviation: KO=knockout. Scale bars: C-E=11μm; G,H=500nm.

### ARSI is a novel sulfatase of 4-chondroitin sulfate (C4S)

To determine the biochemical activity of the orphan sulfatase ARSI *in vitro*, human *ARSI* was overexpressed in HeLa cells. RT-qPCR showed that HeLa cell clones transfected with an empty vector expressed *ARSI* at very low levels, while *ARSI* was successfully overexpressed in *ARSI*-transfected HeLa cell clones (Fig. 6A). HPLC analyses of harvested CS disaccharides demonstrated that total CS amounts were significantly reduced in stable *ARSI*-transfected HeLa clones, compared to stable empty vector-transfected control clones (Fig. 6B). *ARSI* expression caused the ratio of C4S to total CS to significantly decrease, while the ratio of non-sulfated C0S to total CS significantly increased (Fig. 6C). Arguing against any complication from genetic compensation in this overexpression experiment, *ARSI*-transfected HeLa clones did not alter the expression of genes that promote CS sulfation, such as the sulfotransferases *CHST3*, *CHST11*, and *CHST12*, or the glucuronyltransferase *B3GAT3*, compared to empty vector-transfected HeLa clones (Fig. 6D).

**Figure 6.**
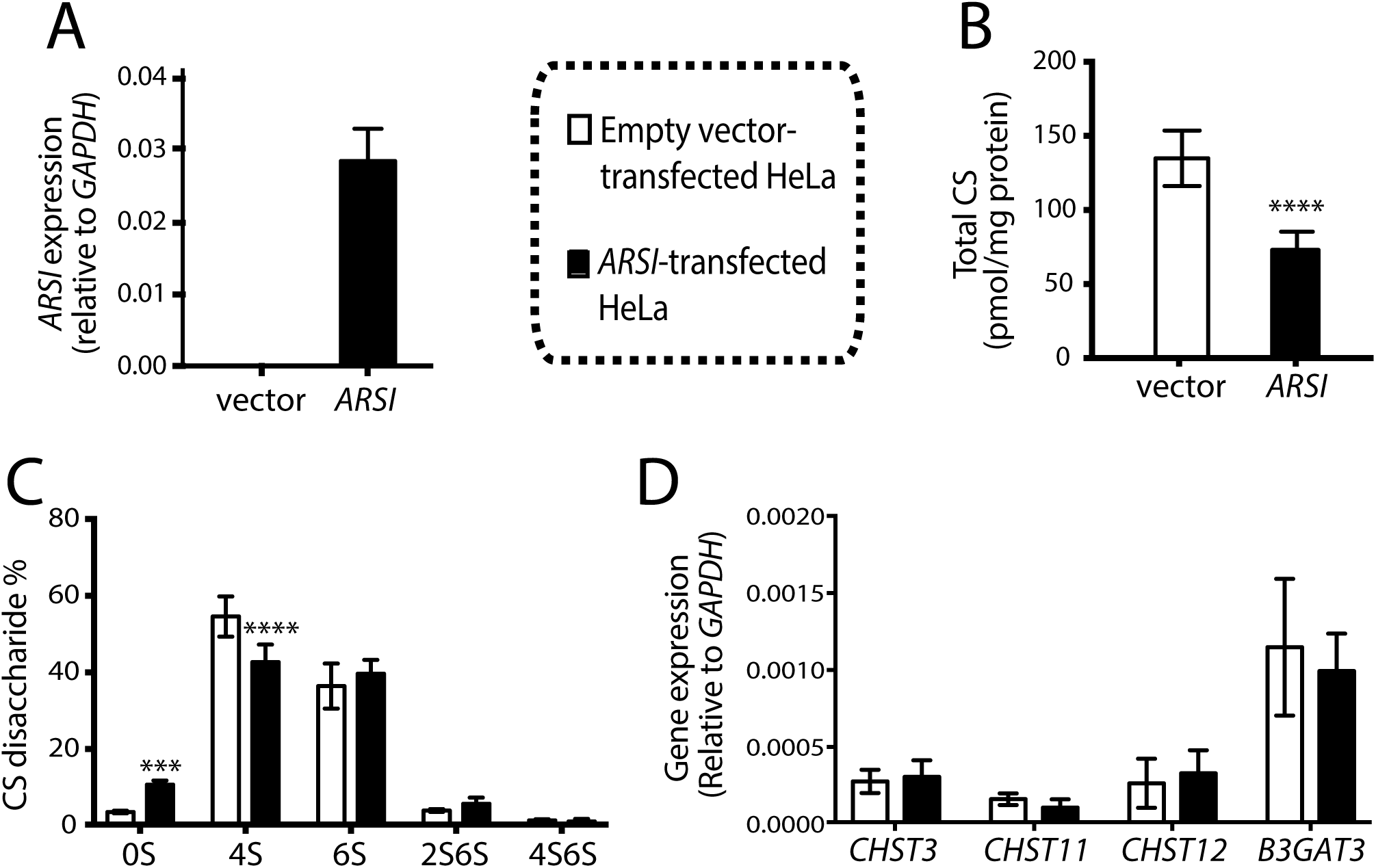
*ARSI* overexpression in HeLa cells reduced sulfation of CS GAGs. **(A)** RT-qPCR showed that human *ARSI* was successfully overexpressed in transfected HeLa cells. **(B)** HPLC demonstrated a decrease in total CS in *ARSI*-transfected HeLa clones, compared with empty vector controls. **(C)** The ratio of 4S to total CS decreased, while the ratio of non-sulfated 0S increased. **(D)** RT-qPCR revealed that the expression of enzymes that promote CS sulfation, like *CHST3, CHST11,* and *CHST12,*and the glucuronyltransferase *B3GAT3* were not affected. ***p<0.0002; ****p<0.0001. Abbreviations: 0S=non sulfated chondroitin, 2S=2-sulfated chondroitin, 4S=4-sulfated chondroitin, 6S=6-sulfated chondroitin.

To further evaluate ARSI sulfatase activity, RCS *Arsi^-/-^* chondrocyte clones underwent HPLC analyses. The percentage of C4S significantly increased in RCS *Arsi^-/-^* chondrocyte clones (Fig. 7), producing a significant increase of the overall charge density from 0.79 to 0.88. Meanwhile, the non-sulfated C0S significantly decreased, and no changes to the other disaccharides sulfated in different positions were observed (Fig. 7).

**Figure 7.**
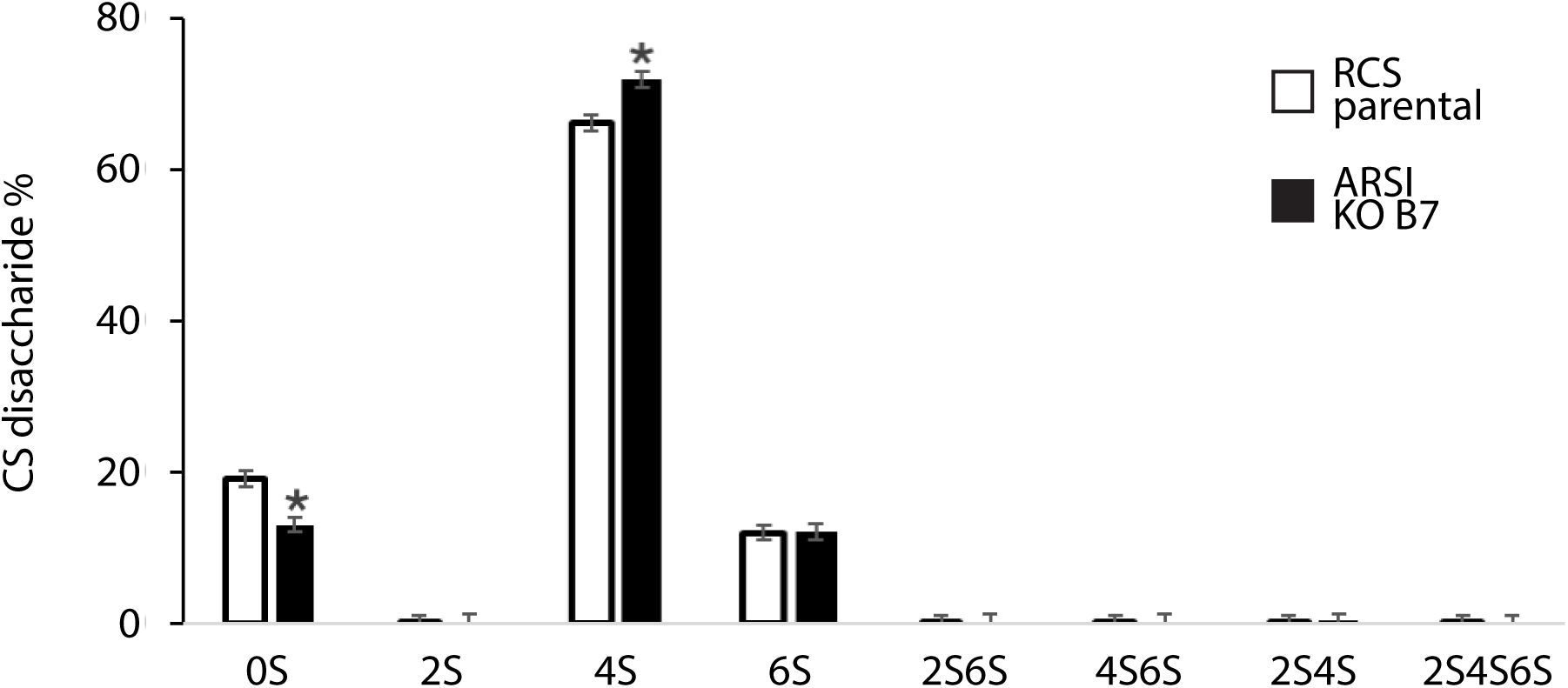
ARSI knockout in RCS chondrocytes increased sulfation of C4S. HPLC analyses demonstrated that the loss of ARSI in RCS chondrocytes caused the percentage of C4S to significantly increase, while the percentage of non-sulfated CS significantly decreased. *p<0.05. Abbreviation: KO=knockout.

To estimate specific activity of ARSI on CS sugars, human *ARSI* was purified biochemically and incubated with GalNAc4S and GalNAc6S. A twelve-hour incubation of GalNAc4S in 25 mM sodium acetate (pH 4.5) with 410 ng of protein A-fused ARSI expressed in the presence of SUMF1 gave rise to 0.5 nmol of GalNAc (Fig. 8). Thus, the specific activity on GalNAc4S of protein A-fused ARSI was estimated to be 0.1 nmol/μg/hr. No activity of ARSI was detected on GalNAc6S in this assay (Fig. 8). In total, these gain- and loss-of-function biochemical data revealed that ARSI is a novel chondroitin sulfatase that acts on GalNAc4S at the nonreducing terminal of the CS/DS chain.

**Figure 8.**
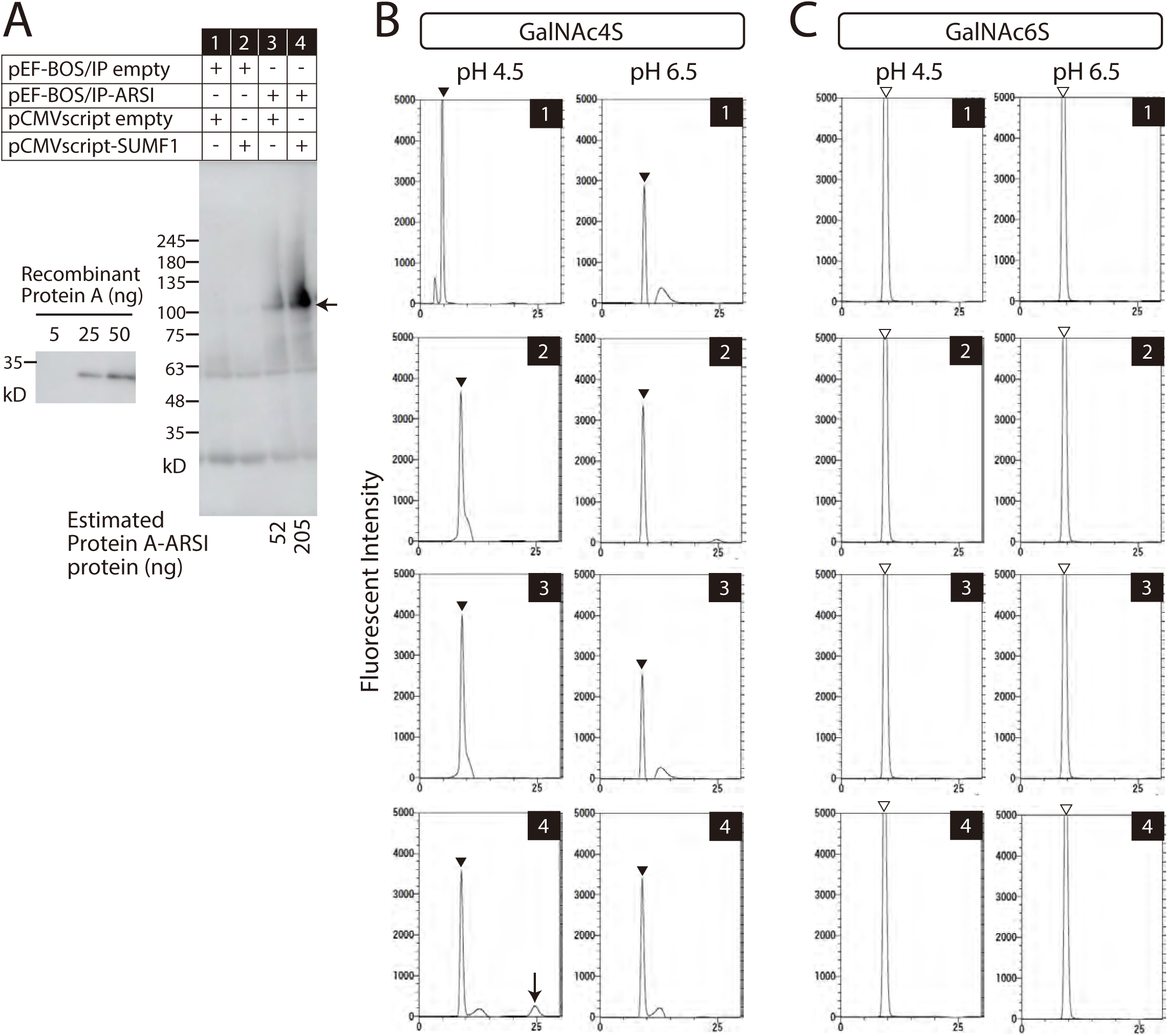
Specific activity of ARSI on GalNAc4S was 0.1nmol/μg/hr. **(A)** COS-7 cells were transfected with (+) or without (-) pEF-BOS/IP empty vector, pEF-BOS/IP-ARSI, pCMVscript empty vector, or pCMVscript-SUMF1. Western blotting showed that more Protein A-fused ARSI (arrow) was secreted into the culture media when ARSI and SUMF1 were co-transfected (lane 4). Using known concentrations of recombinant Protein A, the amount of Protein A-fused ARSI was calculated. **(B)** The same groups of transfected COS-7 cells (Lanes 1-4, panel A) were used as enzyme sources for HPLC analyses of GalNAc4S as a substrate for desulfation at pH4.5 or pH6.5. The filled arrowheads indicate the elution position of GalNac4S. Only enzymes obtained from cells transfected with pEF-BOS/IP-ARSI and pCMVscript-SUMF1 caused production of the desulfated GalNAc at pH4.5 (arrow, #4). **(C)** Identical experiments using GalNAc6S as a substrate did not show activity. The open arrowheads indicate the elution position of GalNac6S.

### ARSI inhibited chondrocyte maturation *in vitro*

RCS chondrocytes can model chondrocyte maturation *in vitro* by treatment with fibroblast growth factor (FGF; Kurimchak et al., 2013). To evaluate if ARSI regulates chondrocyte maturation, RCS *Arsi^-/-^* chondrocytes were stimulated to undergo maturation by addition of FGF18, and maturation markers were analyzed by RT-qPCR. Right before FGF18 treatment, *Arsi^-/-^* chondrocytes already had significantly increased expression of the maturation markers *Mmp13* and *Vegfa*, compared to parental chondrocytes (Fig. 9). They also had significantly increased *Col2a1* expression. In parental RCS chondrocytes, FGF18 treatment for 24 hours caused significant increases in the maturation markers *Mmp13* and *Vegfa*, compared to parental cells before treatment, verifying FGF18 induced chondrocyte maturation (Fig. 9). In *Arsi^-/-^* chondrocytes, the maturation marker *Mmp13* significantly increased after FGF18 treatment, compared to *Arsi^-/-^* chondrocytes before FGF18 addition (Fig. 9). *Col2a1* is known to be down-regulated during chondrocyte maturation (Eames and Helms, 2004), and *Arsi^-/-^* chondrocytes had significantly decreased expression of *Col2a1* after FGF18 treatment.

**Figure 9.**
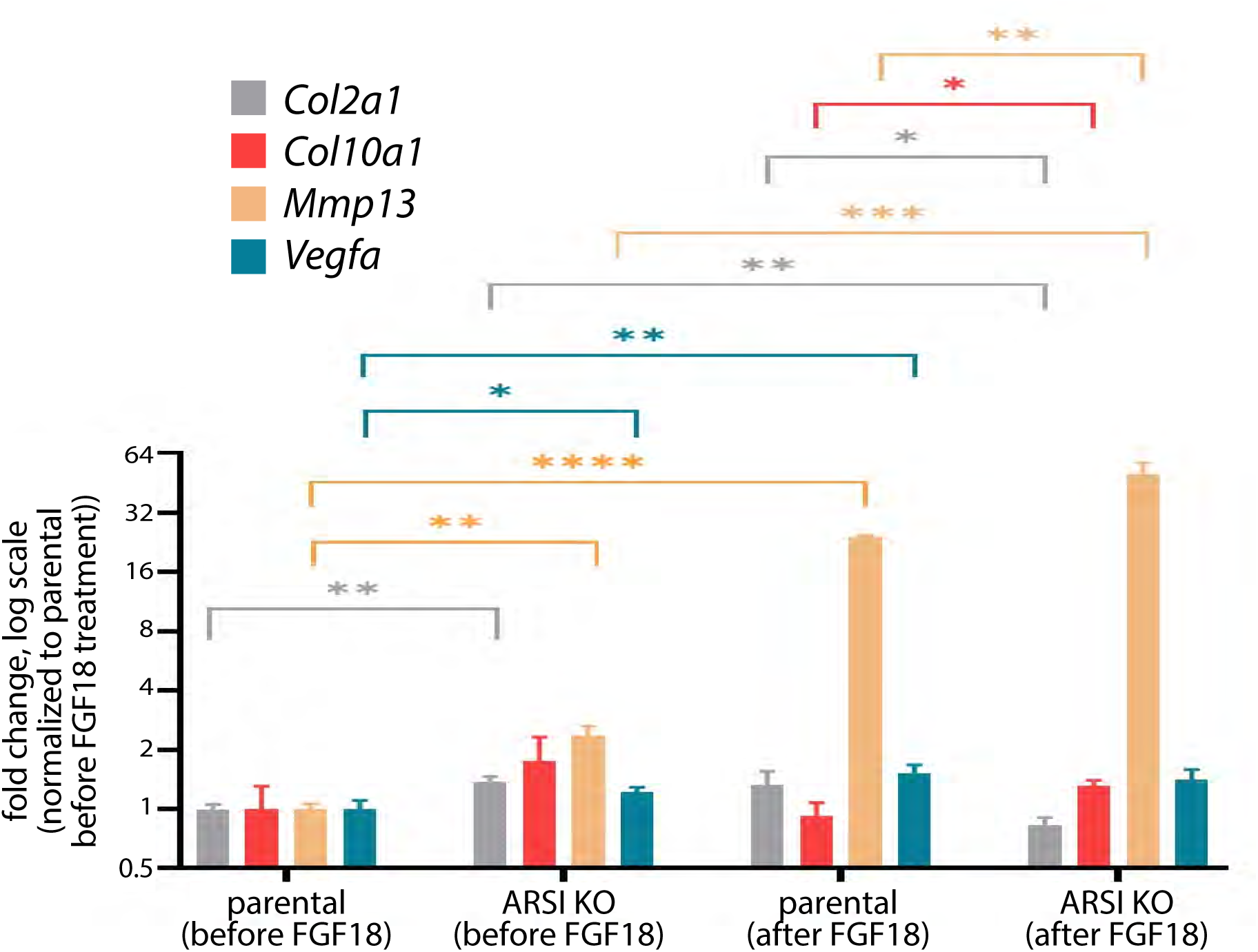
ARSI inhibited chondrocyte maturation *in vitro*. RT-qPCR revealed that *Arsi^-/-^* RCS chondrocytes had significantly increased expression of *Col2a1* and the maturation markers *Vegfa* and *Mmp13*, compared to parental chondrocytes (n=3). After induction of maturation by FGF18 for 24 hours, *Arsi^-/-^* chondrocytes expressed the maturation markers *Col10a1* and *Mmp13* significantly higher than FGF-treated parental chondrocytes, also with significantly decreased *Col2a1* (n=3). *p<0.05; **p<0.01; ***p<0.001; ****p<0.0001. Abbreviations: KO=knockout.

Comparing parental and *Arsi^-/-^* chondrocytes after FGF18 induction of maturation, important differences were observed. *Arsi^-/-^* chondrocytes expressed the maturation markers *Col10a1* and *Mmp13* significantly higher than FGF-treated parental chondrocytes (Fig. 9). Also, *Arsi^-/-^* chondrocytes had significantly decreased expression of *Col2a1* after FGF18 treatment, compared to parental RCS cells (Fig. 9). These *in vitro* data showed that ARSI normally inhibits chondrocyte maturation, since maturation markers were significantly increased in *Arsi^-/-^* chondrocytes.

## Discussion

Modifications to the ECM of the developing growth plate during endochondral ossification has focused on collagens (Ortega et al., 2003), but similar changes to PGs have not been fully characterized (Deutsch et al., 1995; Farquharson et al., 1994; Hackett et al., 2016; Wuthier, 1969). Here, we used synchrotron imaging to illustrate for the first time that *O*-linked sulfate esters, reflecting PG sulfation, decreased significantly during cartilage maturation in developing long bones. Interestingly, the orphan sulfatase ARSI was differentially expressed in this same region, while the known chondroitin sulfatases ARSB and GALNS were not. Following this lead, we revealed the first biochemical characterization of ARSI as a lysosomal CSPG sulfatase, specifically targetting GalNAc4S on the nonreducing terminal of CS/DS. Our results supplement the scarce literature about ARSI and potentiate future work uncovering its role in skeletal development and disease.

*ARSI* was identified decades ago as an arylsulfatase through bioinformatic analysis, but many questions about its biochemical and biological roles remain to be unveiled (Sardiello et al., 2005). General sulfatase activity of ARSI was described using the artificial substrate 4-methylumbelliferyl sulfate (Oshikawa et al., 2009), but its biological substrate remained elusive. ARSI might have been predicted to function like ARSB, whose substrate is GalNAc4S, because they have high sequence similarity (Obaya, 2006; Sardiello et al., 2005). Indeed, experiments here confirmed that ARSI is a novel CSPG sulfatase. Overexpression of ARSI in HeLa cells significantly decreased the relative levels of GalNAc4S, while knockout of ARSI in RCS chondrocytes significantly increased relative levels of GalNAc4S. Importantly, in a cell-free system, ARSI showed specific activity on GalNAc4S (0.1 nmol/μg/hr), but not on GalNAc6S.

Why have two C4S sulfatases? Spatial restriction of activity might answer this question. As an example, SULF1 and SULF2 have redundant biochemical specificity for removing 6-*O* sulfates from heparan sulfate PGs (Diez-Roux and Ballabio, 2005). However, their expression patterns in the body are different, and thus, loss of *Sulf1* has a different phenotype from loss of *Sulf2* (Kalus et al., 2009). Similarly, ARSB and ARSI might both have specificity for C4S, but our extensive expression analyses (i.e., RNA sequencing, RNA *in situ* hybridization, and immunofluorescence) showed that their expression patterns are different. In developing cartilage of both mouse (E14.5) and chick (HH36), ARSI, but not ARSB, was specifically expressed in mature cartilage, expanding upon a previous report (Ratzka et al., 2010). Mouse genetic models suggest that some *Arsb^-/-^* bone phenotypes can be rescued by enzyme replacement therapies in osteoclasts and osteocytes, not chondrocytes (Hendrickx et al., 2020; Pohl et al., 2018). Considering ARSI’s specific expression in mature cartilage, it will be interesting to elucidate how the *Arsb^-/-^* phenotype compares with any skeletal defects in *Arsi^-/-^* models.

Our ARSI data highlight that the relationship between PG sulfatases and lysosomes is both biochemical and cell biological. Many PG sulfatases, such as ARSB, GALNS, and IDS, reside within lysosomes, where they participate in the stepwise degradation of glycosaminoglycans (GAGs; Brown and Eames, 2016). Our colocalization studies, using immunofluorescence and lysosomal markers, strongly suggest that ARSI also localizes to lysosomes, where it may function. Despite a report that ARSI did not colocalize with the lysosomal marker cathepsin D in the retinal-derived cell line ARPE-19 (Oshikawa et al., 2009), EGFP-tagged ARSI colocalized with both LAMP1-positive lysosomes and RAB5-positive endosomes in HeLa cells, suggesting that it plays a role in the autophagy lysosomal pathway. From a biochemical perspective, lysosomal PG sulfatases show optimal activity in the acidic environment of lysosomes (Diez-Roux and Ballabio, 2005). Further supporting its identity as a lysosomal PG sulfatase, ARSI showed C4S activity at pH of 4.5, but not at pH of 6.5, in our isolated cell-free system. This stands in contrast to a report showing extracellular ARSI activity at neutral conditions in ARPE-19 cells (Oshikawa et al., 2009), but activity in that study was measured on an artificial substrate, and endogenous arylsulfatases of ARPE-19 show maximal activity at pH5-6. Future work could test whether ARSI can act as an endo-type sulfatase in neutral conditions. In humans, mutations to some PG sulfatases can cause various forms of mucopolysaccharidosis (MPS), a group of inherited lysosomal storage disorders (LSD’s) characterized by the accumulation of partially-degraded GAGs within lysosomes (Diez-Roux and Ballabio, 2005). Altered lysosome homeostasis is a diagnostic feature of LSD’s, including MPS’s (Bajaj et al., 2019; Settembre and Perera, 2024), presumably as a compensatory mechanism for decreased lysosome-mediated catabolism. In this context, our findings of increased lysosome numbers and sizes in *Arsi^-/-^* chondrocytes underscore the importance of establishing and evaluating *Arsi^-/-^* animals as potential models of LSD’s.

Our chemically-specific XRF imaging approach revealed that PG sulfation decreases during cartilage maturation. Previous studies using biochemical and histological methods suggested a reduction of C4S and C6S in mature cartilage of growth plates in fetal calves and juvenile chicks (Farquharson et al., 1994; Wuthier, 1969). We confirmed that the sulfur signal in cartilage is comprised mostly of *O*-linked sulfate esters, which is the form of sulfur in sulfated PGs (Mikami and Kitagawa, 2013). Indeed, we previously showed that cartilage of PG-deficient *fam20b* mutant zebrafish had reduced *O*-linked sulfate esters by XRF imaging (Hackett et al., 2016). We also illustrated for the first time the exact spatial distribution of PG sulfation across an intact growth plate. Interestingly, decreased PG sulfation in mature cartilage of the growth plate does not match previous studies showing higher sulfur levels in mature cartilage of articular cartilage in horses and pigs using X-ray emission techniques (Reinert et al., 2001; Rizzo et al., 1995), suggesting differences in PG homeostasis of the transient growth plate and permanent articular cartilage.

Decreasing PG sulfation and ARSI expression during cartilage maturation have implications for understanding molecular mechanisms of endochondral ossification. As mentioned, PG desulfation occurs during PG degradation, so the decrease in PG sulfation might simply reflect ECM degradation during cartilage maturation, similar to collagen degradation by matrix metalloproteinase 13 (Ortega et al., 2003). Our FTIR data did not demonstrate significant PG degradation, however, and suggest another mechanism. Chondrocyte hypertrophy and perichondral bone formation were altered in animal models with reduced PG sulfation, such as brachymorphic (*bm*) and cartilage-matrix-deficient (*cmd*) mice; nanomelic (*nm*) chick; and f*am20b* and *xylt1* mutant zebrafish (Cortes et al., 2009; Domowicz et al., 2009; Eames et al., 2011; Watanabe et al., 1994). Sulfated PGs are known to bind growth factors and influence growth factor signalling, and aberrant signalling in many growth factor pathways has been described in developing cartilage of PG-defective animal models (Cortes et al., 2009; Kluppel et al., 2005; Koosha et al., 2024). Therefore, decreased PG sulfation during normal cartilage maturation might release many growth factors, impacting several signalling pathways that can regulate multiple aspects of endochondral ossification. Indeed, a functional role for ARSI in endochondral ossification is supported by our *in vitro* findings of increased maturation markers in *Arsi^-/-^* chondrocytes. Taken together, our data suggest that ARSI might cause a decrease in PG sulfation during cartilage maturation, potentially altering growth factor-dependent aspects of endochondral ossification. Therefore, the establishment and analyses of *Arsi^-/-^* animal models is an important future direction of research, and *ARSI* mutations should be considered for human skeletal diseases of unknown etiology.

## Materials and methods

### Animal tissues

All animal experiments were approved by the University of Saskatchewan’s Animal Research Ethics Board and adhered to the Canadian Council on Animal Care guidelines for humane use. White leghorn chicken eggs (University of Saskatchewan) were incubated in a humidified, rocking incubator at 37°C, and C57/BL6 mouse embryos were collected from pregnant dams (University of Saskatchewan). Tissues were processed as previously described for histological and gene expression studies (Gomez-Picos et al., 2025), unless otherwise detailed below, or for XRF imaging (Hackett et al., 2016). For FTIR analysis, sections were melted onto an IR-grade CaF2 window (Crystran).

### Histology

Chick limbs were subjected to acid-free Alizarin red and Alcian blue whole-mount staining, as previously described (Eames et al., 2011). Chick humerus sections were stained with Safranin O, as described (McManus and Mowry, 1960), but the Safranin O staining step was only 20 min for sample in Fig. 1C.

### Section immunofluorescence

For unfixed sections adjacent to those imaged at the synchrotron, slides were dried at 60°C for 30 min, fixed in 4% PFA overnight at 4°C, and rinsed twice in PBST (PBS/0.5% triton-100; Fisher Scientific). Alternatively, fixed sections were dried for 10 min, post-fixed in 4% PFA at room temperature (RT) for 20 min, and rinsed twice in PBST. Afterwards, section immunos were performed as described (Eames and Schneider, 2005). Antibodies were: anti-COLX (X-AC9, Developmental Studies Hybridoma Bank (DSHB); 1:100); anti-COL2 (II-II6B3, DSHB; 1:100); anti-ARSI (SAB4501497, Sigma; 1:300); anti-GALNS (ab231647, Abcam; 1:100); anti-ARSB (ab85727, Abcam; 1:300); Alexa Fluor Plus 488 goat anti-rabbit IgG (H+L) (A32731, Thermo Fisher); Alexa Fluor Plus 488 goat anti-mouse IgG (H+L) (A32723, Thermo Fisher); Alexa Fluor 594 goat anti-rabbit IgG (H+L) (A11037, Thermo Fisher).

### Synchrotron imaging and analyses

As previously described (Hackett et al., 2016), XRF imaging of total sulfur was performed at the VESPERS beamline, and chemically-specific XRF imaging and XANES measures were performed at the SXRMB beamline of the Canadian Light Source (Saskatoon, SK). Data processing was as described (Hackett et al., 2016) with the following modifications. Immature and mature cartilage ROIs were drawn on the XRF image by importing the COLX immunofluorescence image from the adjacent tissue section in Adobe Photoshop CS6 (Adobe Inc., San Jose, CA) as a new layer, and total pixels and average pixel intensity were counted from the histogram.

### Transcriptomic dataset

The mouse E14.5 RNAseq data are in the Gene Expression Omnibus (GEO) database (NIH) under accession GSE110051.

### Human *ARSI* expression constructs

Each constructed plasmid was confirmed by sequencing the entire coding region and ligation joints. For overexpression experiments, a full-length form of ARSI was amplified from human placental cDNA library using a pair of primers for the first round of nested PCR; 5’ primer (5’-gccggc<u>agatct</u>**atg**cacaccctcactg-3’) containing *Bgl*II site (underlined) and the start codon (bold), and 3’ primer (5’-cagttttctccc<u>agatct</u>ccatc-3’) containing *Bgl*II site (underlined). For second round of amplification, a pair of primers were used; 5’ primer (5’-gccggc<u>agatct</u>**atg**cacaccctcactg-3’) containing *Bgl*II site (underlined) and the start codon (bold), and 3’ primer (5’-tctccc<u>tctaga</u>ccatcagatccgttg-3’) containing *Bgl*II site (underlined). After adding an A-tail to the PCR fragment by Go Taq Flexi polymerase (Promega, Madison, WI), the cloned sequence was subcloned to pGEM-T easy vector (Promega) (pGEM-T ARSI(FL)). For ARSI-EGFP fusion, a 5’-primer (5’-tctcgagctcaagctaccatgcacaccctcact-3’), and a 3’-primer (5’-ggcgaccggtggatcgatccgttgggacattagc-3’) amplified ARSI from pGEM-T ARSI(FL), and this was cloned into pEGFP-N1 vector (Clontech, San Jose, CA), digested with *Hind*III and *BamH*I, based on site-specific recombination using the In-Fusion cloning kit (Clontech), creating pEGFP-N1-ARSI. Additionally, full-length human *ARSI* was cloned into pSP72-EF1-IRES-Blast vector.

For biochemical experiments, a truncated form of ARSI, lacking the first NH2-terminal consisting of 24 amino acids, was amplified from human placental cDNA using a 5’ primer (5’-ccg<u>agatct</u>gtggccgacgggccc-3’) containing an in-frame *Bgl*II site (underlined) and a 3’ primer (5’-tctccc<u>agatct</u>ccacca**tca**gatccgttg-3’) containing an in-frame *Bgl*II site (underlined) and the stop codon (bold). The PCR fragment was digested with *Bgl*II and subcloned into the *BamH*I site of the pEF-BOS/IP vector (Mizushima and Nagata, 1990; Nadanaka et al., 2014), resulting in pEF-BOS/IP-ARSI, where ARSI was fused to the insulin signal and protein A sequences.

### HeLa lines and analyses

For immunolocalization and biochemical studies, pEGFP-N1-ARSI (2μg) and pEF-BOS/IP-ARSI) (2μg), respectively, were transfected into HeLa cells using Lipofectamine 3000 (Thermo Fisher Scientific, Waltham, MA) according to the manufacturer’s instructions. For GAG and RT-qPCR analyses, pSP72-EF1-hARSI-IRES-Blast plasmid was linearized with ClaI and then transfected to HeLa cells using FuGENE6 (Promega).

Transfected cells were selected with blasticidin and multiple clones stably expressing ARSI were established. To obtain negative control cells, pSP72-EF1-IRES-Blast empty vector was transfected to HeLa cells.

For **immunofluorescence**, HeLa cells cultured on a 3.5-cm glass-bottomed dish (AGC Techno Glass Co. Ltd., Shizuoka, Japan) were fixed with phosphate buffered saline (PBS) containing 4% paraformaldehyde for 30 min on ice. After washing with PBS, cells were permeabilized with PBS containing 0.2% Triton X-100 for 10 min at RT. After blocking with PBS containing 2% bovine serum albumin (BSA) for 1 hour at RT, cells were incubated overnight at 4°C with the following antibodies: anti-GFP (M048-3; Medical & Biological Laboratories Co., Tokyo, Japan; 1:10,000), anti-LAMP1 (sc-17768, Santa Cruz Biotechnology, Dallas, TX; 1:100) or anti-Rab5 (sc-10767, Santa Cruz Biotechnology; 1:50). After washing with PBS, Alexa Fluor 488 secondary (Thermo Fisher Scientific; 1:400) or Alexa Fluor 594 secondary (Thermo Fisher Scientific; 1:400) was used for the detection of GFP or for LAMP1 and Rab5, respectively. Hoechst33342 was used as the nuclear counterstain. Images were acquired using a Zeiss LSM 700 confocal laser-scanning system (Carl Zeiss Inc., Oberkochen, Germany) equipped with an inverted Axio Observer Z1 microscope. For **RT-qPCR**, total RNA was extracted using High Pure RNA Isolation Kit (Roche) and reverse transcribed with MMLV reverse transcriptase. Real-time PCR was performed using FastStart Essential DNA Green Master (Roche) and LightCycler 96 System (Roche, Basel, Switzerland). Primers used for real-time PCR were as follows: (*ARSI*) F: 5’-caagggggtcaagttggaga-3’, R: 5’-cagcttctgtggcagtgtca-3’; (*B3GAT3*) F: 5’-cctgcctactatctatgttgttac, R: 5’-accaccaggtgtgtgaa-3’; (*CHST3*) F: 5’-tctgccattggcttgaac-3’, R: 5’-catgcagacatgaaatagcaaac-3’; (*CHST11*) F: 5’-aaacgccagcggaagaa-3’, R: 5’-gggatggcagagtgagtaga-3’; (*CHST12*) F: 5’-gagggaaagttctttgtttaagtg-3’, R: 5’-cggccttaacagccataat-3’; (*GAPDH*) F: 5’-atgggtgtgaaccatgagaagta-3’, R: 5’-ggcagtgatggcatggac-3’. For **GAG analysis**, cells were washed with PBS, collected using a cell scraper and homogenized in cold acetone. The dried powder was digested with actinase E (Kaken Pharmaceutical Co. Ltd., Tokyo, Japan) for 2 days at 55°C. Trichloroacetic acid was added to the digested samples (final 5%), and the soluble fraction was obtained. After extraction with diethyl ether, the aqueous phase was subjected to ethanol precipitation. The precipitate was dissolved in a pyridine-acetate buffer and gel-filtrated on a PD-MiniTrap G-25 column (Cytiva, Marlborough, MA). The flow-through fraction was collected, dried up, and dissolved in MilliQ water. An aliquot of the samples was digested with chondroitinase (ChABC and AC-II, Seikagaku Corp., Tokyo, Japan). Digestions proceeded for 2 hours at 37°C and samples were derivatized with 2-aminobenzamide for 2 hours at 65°C. The derivatized samples were analyzed by high-performance liquid chromatography (CBM-20A, Shimadzu) on a YMC-Pack PA-G column (YMC Co., Ltd., Kyoto, Japan). Identification and quantification of the disaccharides were achieved by comparison with chondroitin sulfate disaccharide standard as described previously (Kitagawa et al., 1995).

### Cell-free biochemical analyses

Two days after transfection of 5μg of pEF-BOS/IP-ARSI into COS-7 cells cultured on a 10-cm dish, 1ml of the culture medium was collected and incubated with 10μl of IgG-Sepharose (Cytiva) for 2 hours at 4°C. The beads recovered by centrifugation were washed with Tris-buffered saline containing 0.1% Tween (TBST). The expression of recombinant Protein A-fused ARSI was quantified by Western blotting using a standard of Protein A (160-26191, Fujifilm WAKO Chemicals, Osaka, Japan) at a known concentration. Desulfation reactions using 4-O-sulfated N-acetylgalactosamine (GalNAc4S; G1054, Dextra, Bangkok, Thailand) or 6-O-sulfated N-acetylgalactosamine (GalNAc6S; sc-295652, Santa Cruz Biotechnology) as an acceptor were co-incubated in reaction mixtures containing the following constituents in a total volume of 50μl; 10μg of GalNAc4S or GalNAc6S, 25mM sodium acetate (pH4.5) or 25mM MES-NaOH (pH6.5), 2mM CaCl2, 2mM MnCl2 and 10μl of the resuspended beads. The mixture was incubated at 37°C for 12 hours and lyophilized in a microcentrifuge tube. A 5μl aliquot each of 0.25M 2-aminobenzamide (2-AB) and 1.0M NaCNBH3, both of which were prepared by dissolving in glacial acetic acid/dimethylformamide mixture (3/7, v/v), were added to the sample, and the mixture was incubated at 65°C for 2 hours. The excess reagent was removed via chloroform extraction. The 2-AB-derivatized monosaccharides were analyzed using an HPLC system LC20AD connected to an RF20Axs fluorometric detector (Shimadzu Corp.). Gel filtration chromatography was performed on a TSK-gel G2500PW column (Tosoh Co., Tokyo, Japan) using 20 mM CH3COONH4 (pH7.5) as the effluent at a flow rate of 1ml/min. The eluates were monitored for fluorescence at excitation and emission wavelengths of 330 and 420 nm, respectively.

### RCS lines and analyses

RCS cells, a Swarm chondrosarcoma chondrocyte line (King and Kimura, 2003), were cultured in DMEM High Glucose, supplemented with 10% Fetal Bovine Serum, and 1% penicillin/streptomycin (Euroclone). ARSI loss-of-function RCS clones were obtained through clustered regularly interspaced short palindromic repeats (CRISPR)/CRISPR-associated protein 9 (Cas9) technology. Briefly, 1×10^6^ RCS chondrocytes were transfected with 5 μg of all-in-one vector (Sigma-Aldrich) containing the sgRNA of interest (B7 clone:_tcgtggtacccttggtcat; A2 clone:_gccatggtatcccacgtcg). 48 hours after transfection, the cells were sorted for eGFP fluorescence using the BD FACSAria (BD Biosciences), grown to confluence, and then selected clones were subjected to PCR analysis followed by Sanger sequencing to identify mutations. For FGF18 treatment, cells were grown in 10% FBS (Gibco). Day1 was one day after cells reached confluency, after which cells were treated with 50 ng/ml of human FGF18 (PeproTech) in 0.1% BSA for 24 hours for Day2 samples.

For **Western blot,** RCS chondrocytes were washed three times with PBS and collected using 1% trypsin. The cell pellets were washed at least three times with PBS. Cells were lysed in RIPA lysis buffer (20mM Tris [pH8.0], 150mM NaCl, 0.1% SDS, 1% NP-40, 0.5% sodium deoxycholate) supplemented with PhosSTOP and EDTA-free protease inhibitor tablets (Roche, Indianapolis, IN, USA), using freeze and thawing method. The soluble fractions were isolated by centrifugation at 18,000g for 20 min at 4°C. Total protein concentration in cellular extracts was measured using the colorimetric BCA protein assay kit (Pierce Chemical Co, Boston, MA, USA). Protein extracts, separated by SDS–PAGE and transferred onto PVDF, were probed with primary antibodies overnight against ARSI (HPA038398, Sigma-Aldrich; 1:300) and Filamin A (4762, Cell Signaling Technology; 1:1000). Proteins of interest were detected with HRP-conjugated goat anti-rabbit IgG antibody (Vector Laboratories; 1:2000) and visualized with the ECL Star Enhanced Chemiluminescent Substrate (Euroclone), according to manufacturer’s protocol. Images were acquired using the ChemiDoc-lt imaging system (UVP). For **Lysotracker experiments**, LysoTracker DND99 (L7528 Thermo Fisher) was incubated at 50nM in dark for 40 min at 37°C. Cells were washed three times with PBS 1× and collected. Red fluorescence was measured by FACS Accuri C6, and 10,000 events were collected. For **immunofluorescence**, RCS chondrocytes were fixed for 10 min in 4% PFA and permeabilized for 1 hour in blocking buffer (0.05% (w/v) saponin, 0.5% (w/v) BSA, 50mM NH4Cl, and 0.02% NaN3 in PBS). Cells were incubated in humid chamber for 2 hours at RT with anti-Lamp1 (ab24170, Abcam; 1:200), washed three times in PBS, incubated for 1 hour with AlexaFluor secondary (1:400), washed again three times in PBS, incubated for 20 min with 1μg/ml Hoechst 33342, and mounted in Mowiol (Sigma). Images were acquired using the LSM 880 confocal microscope equipped with a 63×1.4 numerical aperture oil objective, and image quantifications were performed using ImageJ plugins. For **transmission electron microscopy**, cells were fixed with 1% glutaraldehyde in 0.2M HEPES buffer (pH 7.4) for 30 min at RT and post-fixed as described (Polishchuk et al., 2019). After dehydration, the specimens were embedded in epoxy resin and polymerized at 60°C for 72 hours. Thin 60-nm sections were cut on a Leica EM UC7 microtome. EM images were acquired using a FEI Tecnai-12 electron microscope equipped with a VELETTA CCD digital camera (FEI, Eindhoven, The Netherlands). Morphometric analysis on the size of lysosomes was performed using iTEM software (Olympus SYS, Germany). For **GAG analyses**, medium from cultured cells was centrifuged at 5000 rpm 4°C for 10 min, treated with chondroitinase to release single CS non-sulfated and variously sulfated CS disaccharides, derivatized with a specific fluorochrome, and separated by HPLC-MS as previously published (Volpi et al., 2014). For **RT-qPCR**, total RNA was extracted (Qiagen) from cells, and cDNA was synthesized with RevertAid cDNA synthesis kit (Thermo Fisher). RT-qPCR was performed with SsoAdvanced universal SYBR green supermix (Bio-Rad) using CFX96 rtPCR thermocycler device (Bio-Rad). Primer pairs were (*Cyc*, or *Ppia-ps1*, used as internal control) F: 5′-tggaaaagttgtaccccgga-3′, R: 5′-tacctccagtgccgtctcta-3′; (*Col2a1*) F: 5′-ccgatcccctgcagtacatg-3′, R: 5′-tgctctcgatctggttgttc-3′; (*Col10a1*) F: 5′-agctcacggaaaatgaccag-3′, R: 5′-gttctaagcgggggattagg-3′; (*Vegfa*) F: 5′-gtggaagaagaggcctggta-3′, R: 5′-cacacacacagccaagtctc-3′, as published (Cinque et al., 2025). *Mmp13* primers were designed as F: 5′-accagagaagtgtgacccag-3′, R: 5′-catggttgggaag ttctggc-3′; To determine the relative expression of mRNA, 2-ΔΔCT method was used.

### Statistical analysis

Statistically significant differences were determined by the one sample t-test, two-way ANOVA, unpaired t-test, or paired t-test, depending on the results being analyzed. The results are reported as mean ± standard error of the means. p<0.05 was considered statistically significant.

## Acknowledgements

Funding for this paper came from CIHR project grant 148683 (BFE). This work was supported in part by MEXT KAKENHI Scientific Research (B) 20H03386 and 24K02183 (to H. K.) and (C) 21K06089 (to S. N.), a Grant-in-Aid for Challenging Exploratory Research 22K19392 (to H.K.). We thank Rich Schneider for the chick *COL2A1* and *SPP1* RNA *in situ* probes and Ralph Marcucio for the chick *COL10A1* and *IHH* probes. Research described in this paper was performed at the VESPERS and SXRMB beamlines at the Canadian Light Source, which is supported by the Canada Foundation for Innovation, Natural Sciences and Engineering Research Council of Canada, the University of Saskatchewan, the Government of Saskatchewan, Western Economic Diversification Canada, the National Research Council Canada, and the Canadian Institutes of Health Research.

## Supplemental Data

**Supplemental Figure S1. Measured XANES spectra in the HH36 chick humerus fit very well with five standard sulfur compounds. (A)** Absorption curves of standards for 8 different forms of sulfur, normalized to their respective edge-jump values. **(B)** Aggregate fittings of standard curves from the five most abundant chemical forms of sulfur present in chick cartilage matched very well with the averaged XANES spot scans in mature cartilage (n=4).

**Supplemental Figure S2. FTIR imaging suggested that proteins, PGs, and GAGs may decrease in mature cartilage. (A)** IMM and MAT regions (dotted lines) of HH36 chick humerus sections were assigned based on COLX immunostaining of sections adjacent to those used for FTIR imaging. **(B)** The integrated Amide I peak (1590–1720cm^-1^) in the FTIR map was used to approximate total protein levels. **(C)** Normalizing a PG representative band (985–1140cm^-1^) by the Amide I peak suggested a decrease in mature cartilage. **(D)** A GAG second derivative peak (1374cm^-1^), normalized to the integrated Amide I peak, suggested a decrease in mature cartilage. **(E)** Quantitation (n=7) showed no significant differences between MAT and IMM regions (black outlines in B-D), although all PG bands and the amide I band showed slight decreases in mature cartilage. Abbreviations: IMM=immature cartilage; MAT=mature cartilage. Scale bar=1mm.

**Supplemental Figure S3. Alignment of ARSI protein sequences across vertebrate clades.** Putative catalytic sites (highlighted in pink) and the putative catalytic core (boxed in black) are conserved across all compared species, and the entire putative sulfatase domain is underlined in red. A gray asterisk indicates a single, fully conserved residue; a gray colon indicates similar amino acid properties; and a period indicates weakly similar amino acid properties.

**Supplemental Table S1. Amino acid sequences of full-length ARSI were highly conserved across vertebrate clades.** Numbers reflect % identity. Abbreviations: (a)=Arsia; (b)=Arsib; *Dre*=*Danio rerio*; *Gga=Gallus gallus*; *Hsa*=*Homo sapiens*; *Mmu=Mus musculus*.

**Supplemental Figure S4. Chick *ARSI* mRNA expression was in a similar domain of mature cartilage as *IHH*.** RNA *in situ* hybridization on sections of the HH36 chick humerus showed expression of *ARSI* **(A)** in a similar domain of mature cartilage as the prehypertrophic marker *IHH* **(B)**, but not in more mature domains, such as where *COL10A1* **(C)** or *SPP1* **(D)** were expressed, nor where *COL2A1* **(E)** was downregulated. Expression of *GALNS* **(F)** or *ARSB* **(G)** did not show specific expression in immature or mature cartilage regions. Boxed regions in panels A-G indicate regions shown in panels A’-G’, respectively. Arrowheads in panel A show expression of *ARSI* in superficial regions of the cartilage; asterisks (*) in panels A and A’ indicate where the plane of section is tangential through the superficial region. Scale bars: A-G=500μm; A’-G’=200μm.

**Supplemental Figure S5. Immunostaining demonstrated that ARSI was expressed in more superficial regions of mature cartilage in the HH36 chick humerus.** Expression of ARSI spanned the width of tangential sections **(A)**, near the surface of the cartilage element, but deeper sections **(B,C)** demonstrated that protein expression was more restricted to superficial regions of mature cartilage. Yellow arrows illustrate cross-sectional width of the cartilage element. Scale bars=200μm.

**Supplemental Figure S6.** Immunostaining demonstrated that ARSI was expressed in mouse mature cartilage *in vivo* and *in vitro*. Immunostaining on sections of the E14.5 mouse humerus confirmed *in vivo* expression of COLX **(A)** and ARSI **(B)** protein in developing mature cartilage. Alcian blue staining revealed that micromasses of the mouse chondrocyte line ATDC5 differentiated into cartilage by day 7 **(C)**, day 14 **(D)**, and day 21 **(E)**. Increased COLX immunostaining of micromasses **(F-H)** confirmed maturation of ATDC5 cells over 21 days of culture, and ARSI expression **(I-K)** also appeared in ATDC5 micromasses at day 14 and day 21. Red boxes in panels C-E indicate the regions of micromasses used for immunostaining images in panels F-K. **(L)** Western blot revealed increased ARSI expression during mouse ATDC5 chondrocyte maturation *in vitro*. Scale bars: A,B,F-K=100μm; C-E=500μm.

