## supplemental figures/methods for "Arylsulfatase I is a novel lysosomal chondroitin sulfatase regulating endochondral ossification"

A

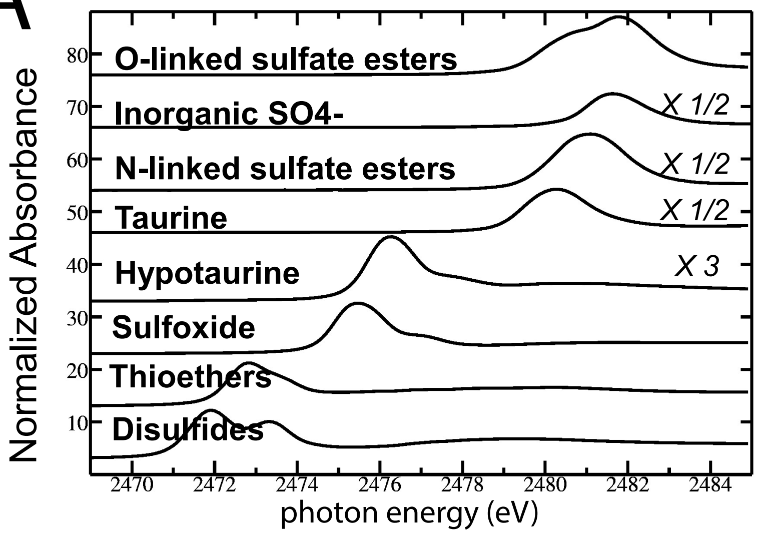


B

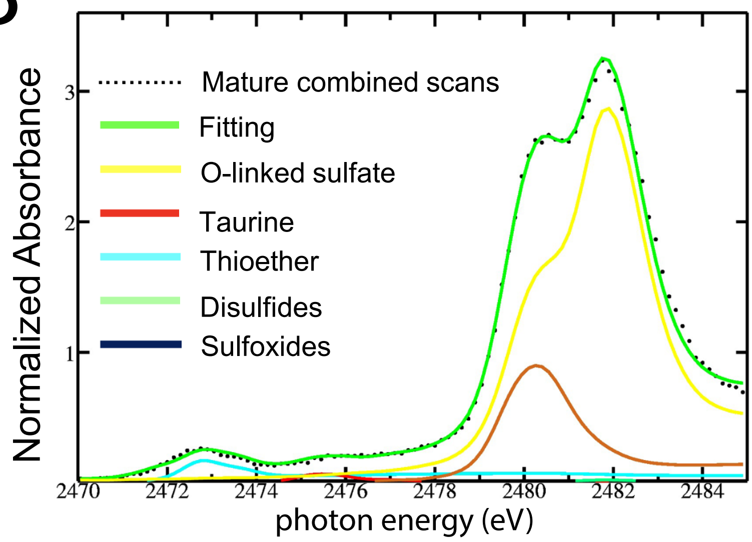


Supplemental Figure 1. Measured XANES spectra in the HH36 chick humerus fit very well with five standard sulfur compounds. (A) Absorption curves of standards for 8 different forms of sulfur, normalized to their respective edge-jump values. (B) Aggregate fittings of standard curves from the five most abundant chemical forms of sulfur present in chick cartilage matched very well with the averaged XANES spot scans in mature cartilage (n=4).

Supplemental Figure 2 Grecco et al.

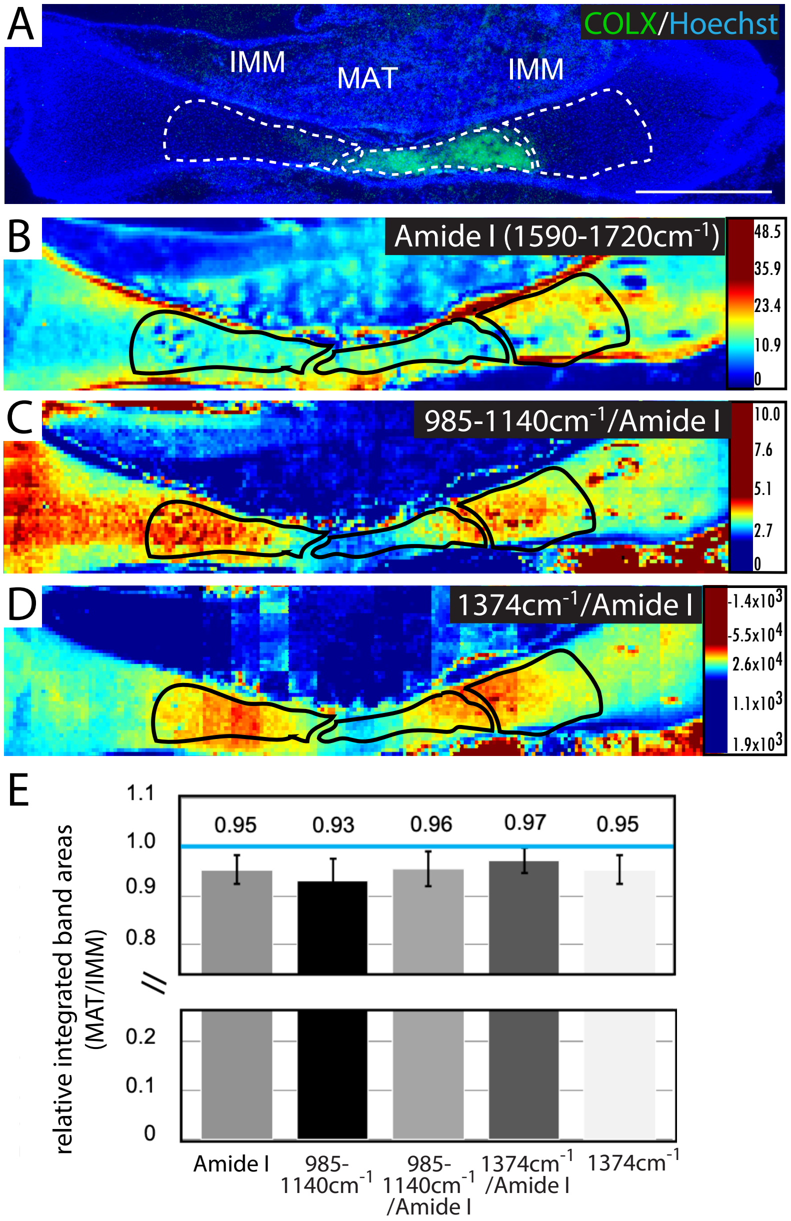


Supplemental Figure 2. FTIR imaging suggested that proteins, PGs, and GAGs may decrease in mature cartilage. (A) IMM and MAT regions (dotted lines) were assigned based on ColX immunostaining of sections adjacent to those used for FTIR imaging. (B) The integrated Amide I peak (1590–1720cm-1) in the FTIR map was used to approximate total protein levels. (C) Normalizing a PG representative band (985–1140cm-1) by the Amide I peak suggested a decrease in mature cartilage. (D) A GAG second derivative peak (1374cm-1), normalized to the integrated Amide I peak, suggested a decrease in mature cartilage. (E) Quantitation (n=7) showed no significant differences between MAT and IMM regions (black outlines in B-D), although all PG bands and the amide I band showed slight decreases in mature cartilage. Abbreviations: IMM=immature cartilage; MAT=mature cartilage. Scale bar=1mm.

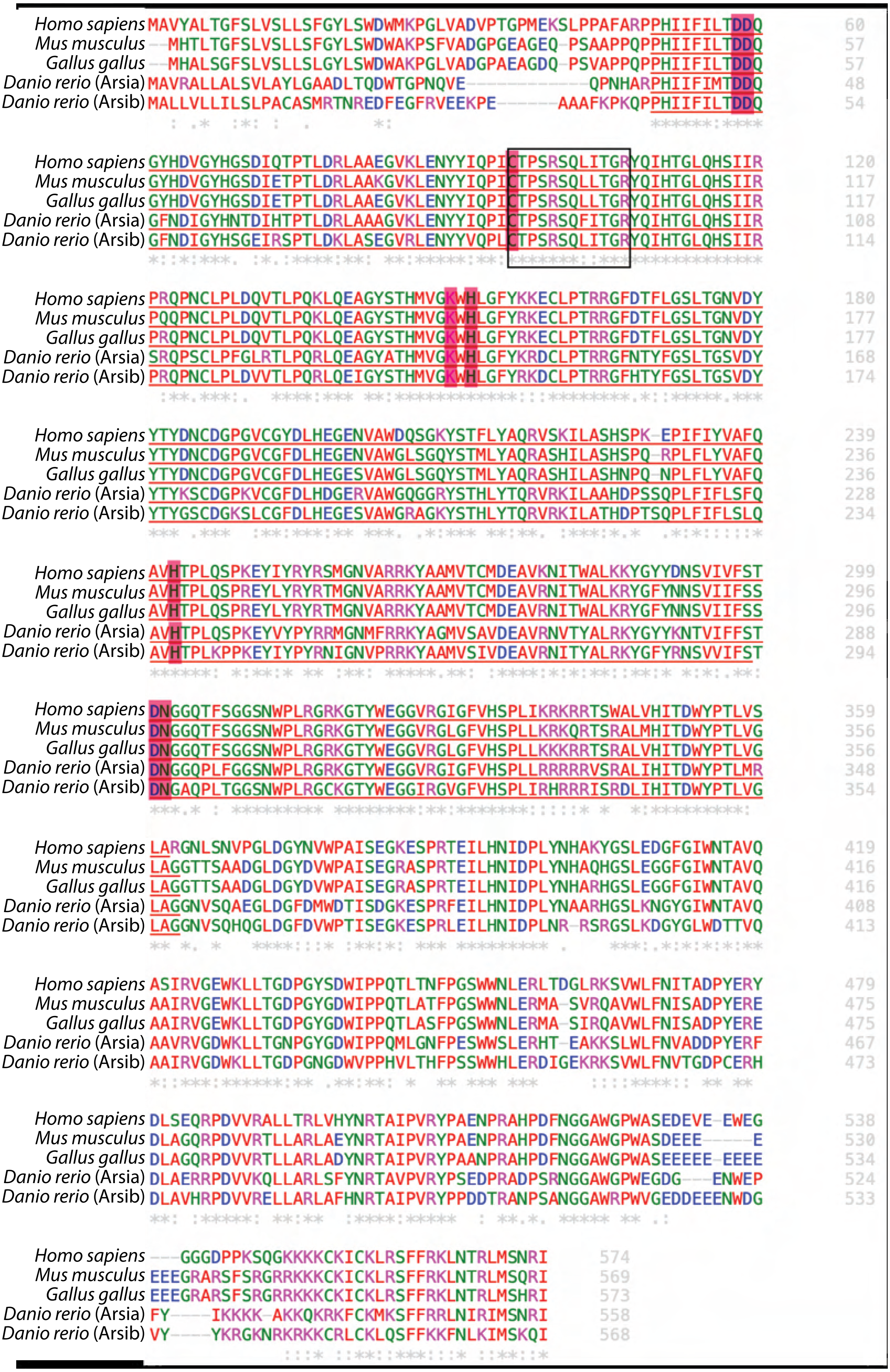


Supplemental Figure 3. Alignment of Arsi protein sequences across vertebrate clades. Putative catalytic sites (highlighted in pink) and the putative catalytic core (boxed in black) are conserved across all compared species, and the entire putative sulfatase domain is underlined in red. A gray asterisk indicates a single, fully conserved residue; a gray colon indicates similar amino acid properties; and a period indicates weakly similar amino acid properties.

Supplemental Table 1. Amino acid sequences of full-length Arsi were highly conserved across vertebrate clades. Numbers reflect % identity. Abbreviations: (a)=Arsia; (b)=Arsib; Dre=Danio rerio; Gga=Gallus gallus; Hsa=Homo sapiens; Mmu=Mus musculus.

| | ***Hsa*** | ***Mmu*** | ***Gga*** | ***Dre* (a)** | ***Dre* (b)** |
|---|---|---|---|---|---|
| ***Hsa*** | 100 | 96.0 | 83.4 | 68.2 | 65.8 |
| ***Mmu*** | | 100 | 83.5 | 68.2 | 66.4 |
| ***Gga*** | | | 100 | 70.6 | 66.7 |
| ***Dre* (a)** | | | | 100 | 71.3 |
| ***Dre* (b)** | | | | | 100 |

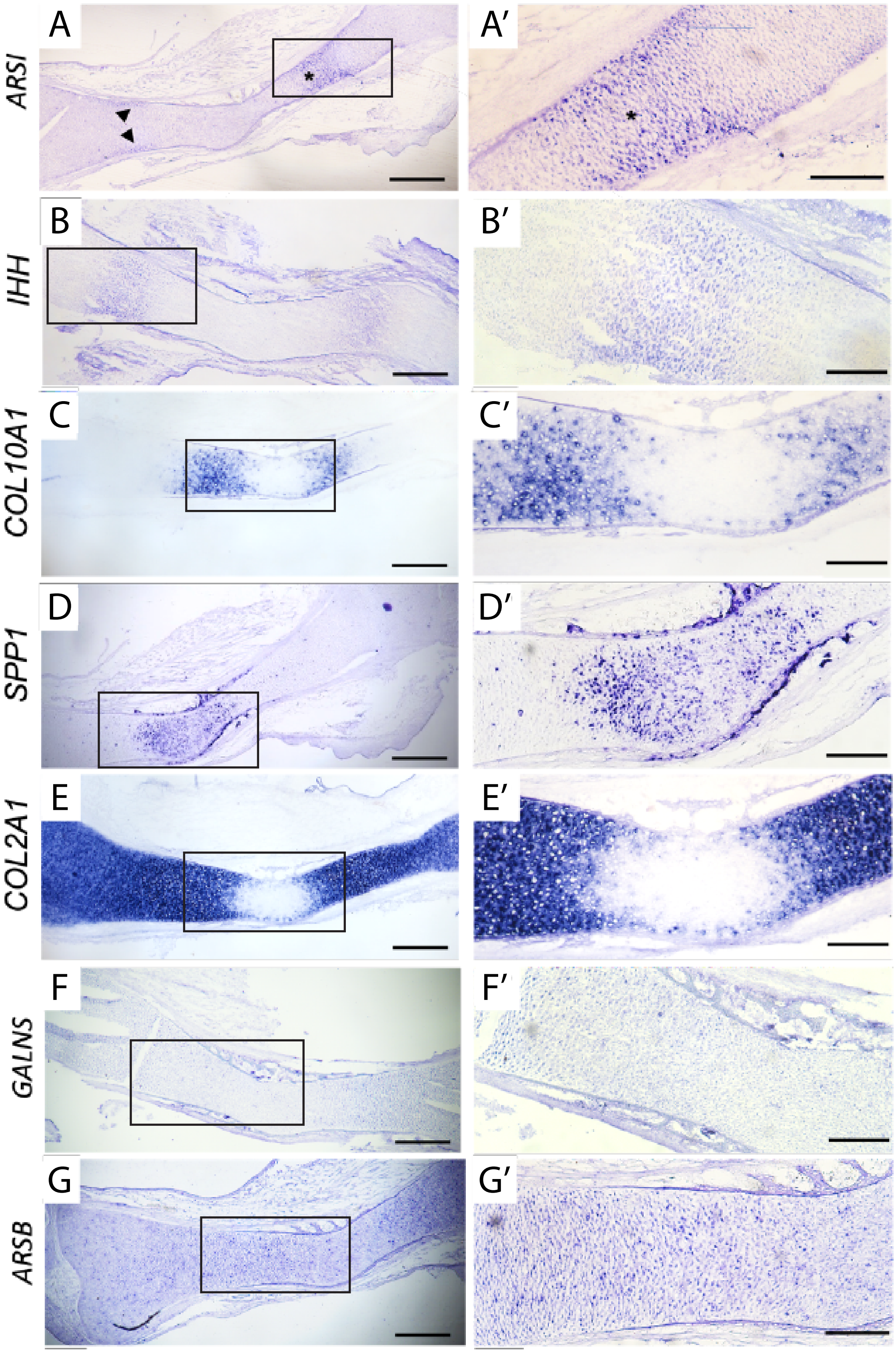


Supplemental Figure 4. Chick ARSI mRNA expression was in a similar domain of mature cartilage as IHH. RNA in situ hybridization on sections of the HH36 chick humerus showed expression of ARSI (A) in a similar domain of mature cartilage as the prehypertrophic marker IHH (B), but not in more mature domains, such as where COL10A1 (C) or SPP1 (D) were expressed, nor where COL2A1 (E) was downregulated. Expression of GALNS (F) or ARSB (G) did not show specific expression in immature or mature cartilage regions. Boxed regions in panels A-G indicate regions shown in panels A′-G′. Arrowheads in panel A and asterisks (*) in panel A′ show expression of ARSI in superficial regions of the cartilage. Scale bars: A-G=500 m; A′-G′=200 m.

### HH36 chick humerus

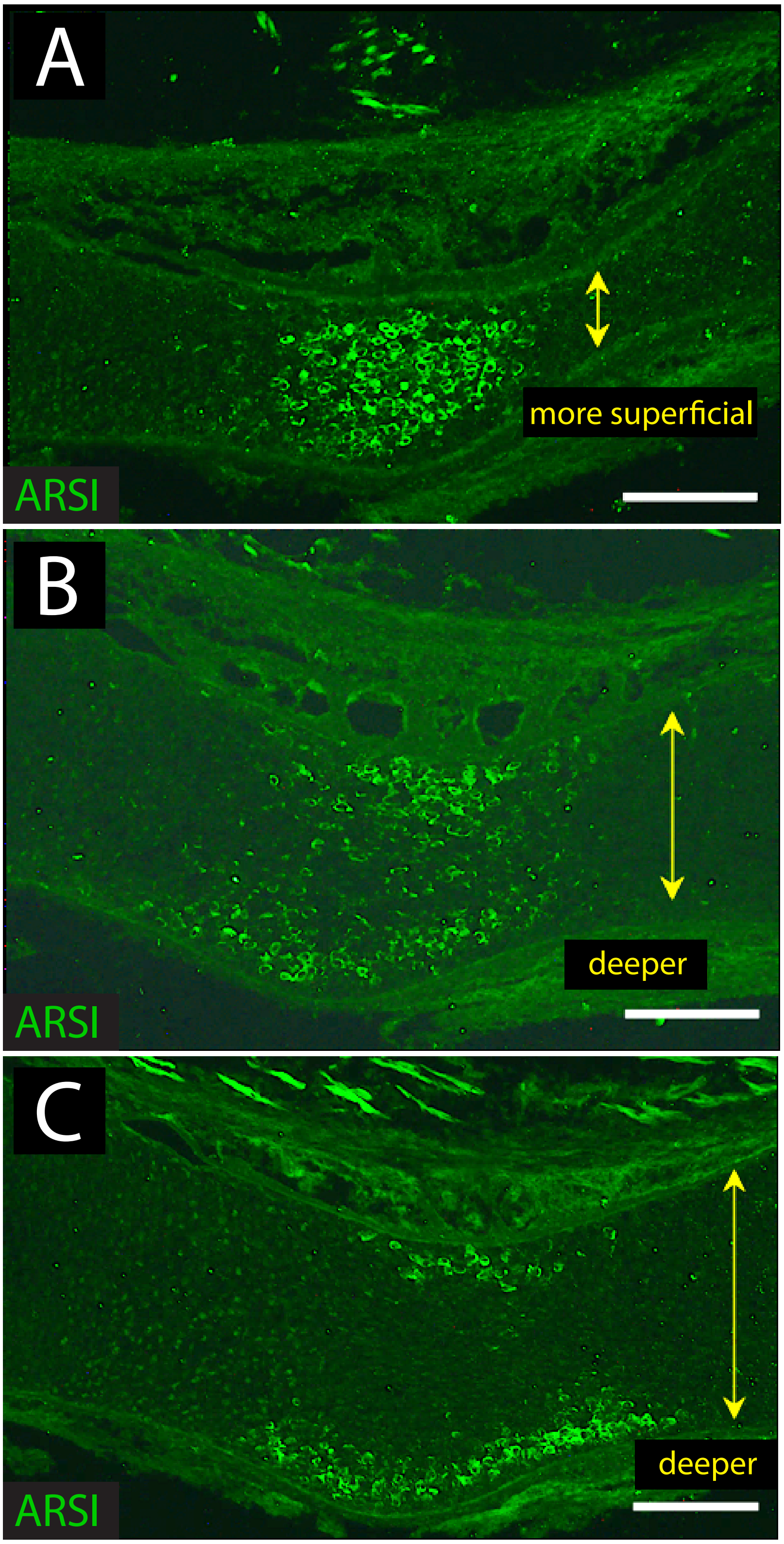


Supplemental Figure S5. Immunostaining demonstrated that ARSI was expressed in more superficial regions of mature cartilage in the HH36 chick humerus. Expression of ARSI spanned the width of tangential sections (A), near the surface of the cartilage element, but deeper sections (B,C) demonstrated that protein expression was more restricted to superficial regions of mature cartilage. Yellow arrows illustrate cross-sectional width of the cartilage element. Scale bars=200  m.

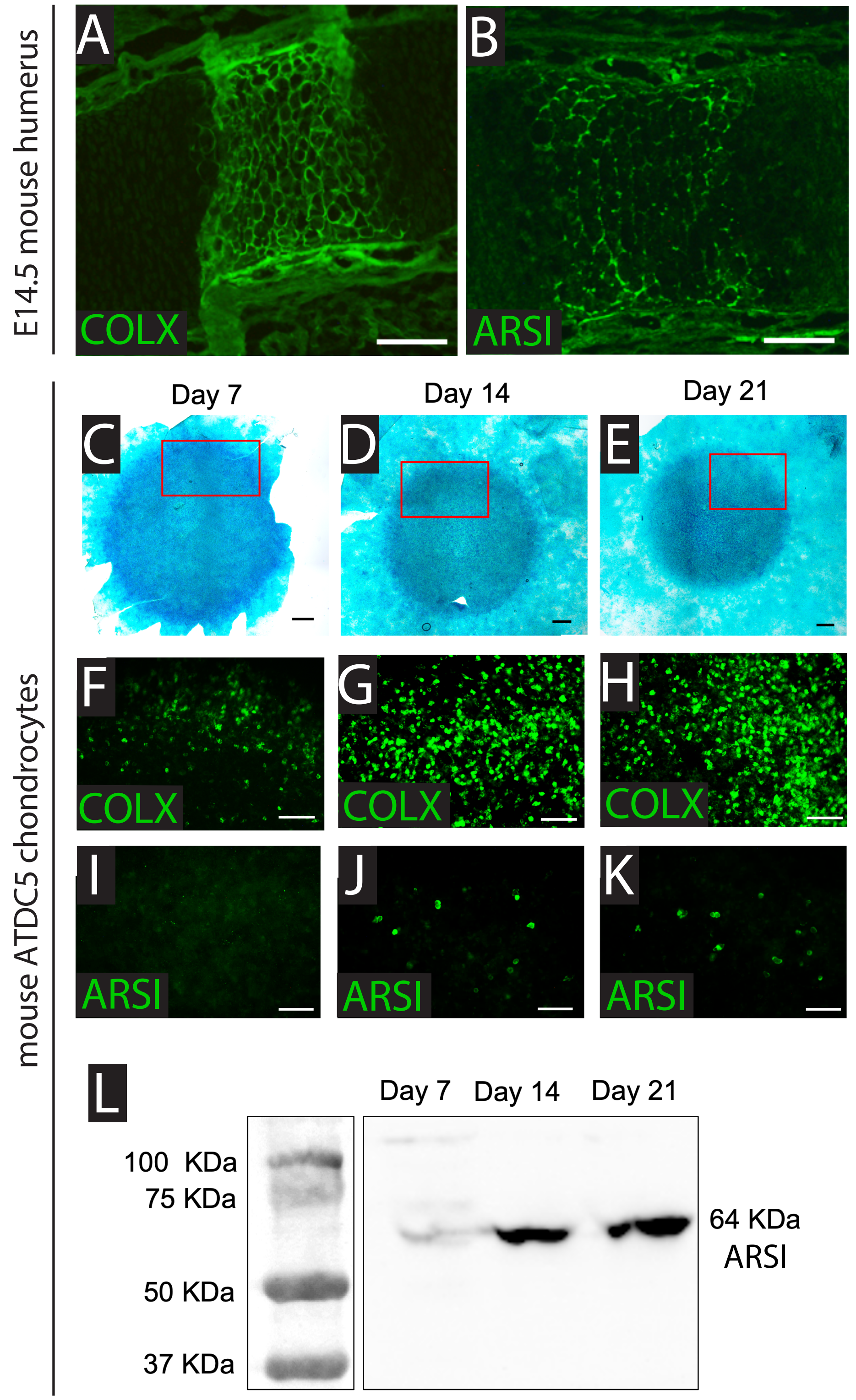


Supplemental Figure S6. Immunostaining demonstrated that Arsi was expressed in mouse mature cartilage in vivo and in vitro. Immunostaining on sections of the E14.5 mouse humerus confirmed in vivo expression of ColX (A) and Arsi (B) protein in developing mature cartilage. Alcian blue staining revealed that micromasses of the mouse chondrocyte line ATDC5 differentiated into cartilage by day 7 (C), day 14 (D), and day 21 (E). Increased ColX immunostaining of micromasses (F-H) confirmed maturation of ATDC5 cells over 21 days of culture, and Arsi expression (I-K) also appeared in ATDC5 micromasses at day 14 and day 21. Red boxes in panels C-E indicate the regions of micromasses used for immunostaining images in panels F-K. Western blot revealed increased Arsi expression during ATDC5 cartilage maturation in vitro. Scale bars: A,B,F-K=100 m; C-E=500 m.

***Fourier transform infrared (FTIR) spectroscopic imaging and analyses***

Data were collected across the spectral region 3600 – 900 $cm^{-1}$ in transmission mode with a Hyperion 3000 FTIR imaging system (Bruker Optics, Ettlingen, Germany) fitted with a mercury-cadmium-telluride (MCT) 64×64 liquid nitrogen-cooled focal plane array (FPA) detector illuminated with a Globar source (aperture 3.5 mm) with a spectral resolution of 4 $cm^{-1}$ with the co-addition of 128 scans, with a background image collected from a blank substrate using 128 co-added scans immediately before each sample. The upper objective of 15x magnification and a numerical aperture of 0.4, combined with a lower condenser of 15x magnification and 0.4 numerical aperture, yielded a pixel size of 21.4 $\mu m^2$, using 8x8 binning. All data processing and image generation was performed using Cytospec software (Cytospec, Version 1.2.04) and Opus software (Version 6.5, Bruker, Ettlingen, Germany). To account for variations in section thickness, normalization of spectra used total protein content using the amide I band (1590 – 1720 $cm^{-1}$; Boskey and Pleshko Camacho, 2007), as described previously (Baker et al., 2014). Three different IR spectral regions of interest were determined in mature versus immature cartilage: the relative abundance of total proteins, and two IR regions representing CS content based on previously validated studies, 985 – 1140 $cm^{-1}$ (Saarakkala and Julkunen, 2010) and the 1374 $cm^{-1}$ second derivative band (Rieppo et al., 2012).

***RNA* in situ *hybridization***

From an HH35 chick forelimb cDNA library, gene fragments were cloned in a pCR™4-TOPO® vector (Thermofisher) for use as *in situ* probes. Primers were: ARSBF: GTAACACGCTGTGCTCTGGA; ARSBR: GGTGGAGGGAACCAATGACC; ARSIF1:

AGGACGAGGTGGAGGAGTG; ARSIR1: AAGCCAGCACAGAGGAAAAG; GALNSF: TCATGGACGACATGGGTTGG; GALNSR: GTGGTTTGCTTGCCACAGAG. Digoxigenin-labelled RNA probes were generated, and *in situ*'s were carried out on chick sections as published (Gomez-Picos et al., 2025). In some cases, probe hydrolysis was done by incubating the probe with twice the volume of RNase-free hydrolysis buffer (40mM NaHCO3/60nM Na2CO3) at 58°C for 10 mins, followed by re-precipitation and dissolution in 0.1% DEPC (Ferrandiz and Sessions, 2008).

***ATCD5 micromass experiments***

ATDC5 cells (RIKEN cell bank) were established into micromasses, and then either stained whole-mount with Alcian blue or immunostained with antibodies in "Immunofluorescence" Methods section, as previously published (Gomez-Picos et al., 2025). For western blot analysis, cells were washed three times with PBS and then dissociated in Laemmli buffer after the addition of proteinase inhibitor (Thermo Fisher). Lysates were centrifuged at 12,000 rpm for 10 min at 4°C, and the supernatant was used for the experiments. Protein extracts were boiled for 5 minutes at 95°C before being loaded onto 10% SDS-PAGE gels and transferred onto nitrocellulose membranes. After blocking with 5% skim milk in PBST (PBS/0.05%Tween-20), the membranes were incubated with anti-ARSI antibody (Sigma) or anti-GAPDH (Sigma) antibodies at 4°C with gentle shaking overnight. Next, the membranes were washed 3 times with 5% skim milk in PBST while shaking at room temperature and incubated with a secondary antibody for 1 h. Membranes were then washed 3 times with 5% skim milk in PBST, 1 time with PBST, and 1 time with PBS. Finally, the chemiluminescent signals on the membranes were detected using an ECL reagent (Thermo Fisher) and a ChemiDoc MP Imaging System (Biorad).
